# Neural latent representation of implicit perceptual decision confidence signatures from brain-wide intracranial EEG

**DOI:** 10.64898/2026.09.30.755781

**Authors:** Umesh Kumar Naik Mudavath, Abdoreza Asadpour, Sabina Gherman, Redmond G. O’Connell, Simon P. Kelly, Stephan Bickel, Shaik Rafi Ahamed, KongFatt Wong-Lin

**Affiliations:** Department of Electronics and Electrical Engineering, Indian Institute of Technology Guwahati, Guwahati, Assam, India.; Intelligent Systems Research Centre, School of Computing, Engineering and Intelligent Systems, Ulster University, Derry∼Londonderry campus, Derry∼Londonderry, Northern Ireland, UK.; School of Life Sciences, University of Sussex, UK.; Feinstein Institutes for Medical Research, Northwell Health, Manhasset, NY, USA.; Trinity College Institute of Neuroscience, and School of Psychology, Trinity College Dublin, Dublin, Ireland.; School of Electrical and Electronic Engineering, and UCD Centre for Biomedical Engineering, University College Dublin, Dublin, Ireland.; Departments of Neurology and Neurosurgery, Zucker School of Medicine at Hofstra/Northwell, Hempstead, NY, USA.; Center for Biomedical Imaging and Neuromodulation, Nathan Kline Institute, Orangeburg, NY, USA.

**Author notes:** Corresponding authors: Umesh Kumar Naik Mudavath; KongFatt Wong-Lin.

**Keywords:** Perceptual decision-making, implicit confidence, stereo electroencephalography sEEG, neural latent embedding, feature selection, convolutional autoencoder, explainable AI

## Abstract

Human perceptual decision confidence has been associated with brain-wide signalling, albeit with inconsistencies across studies. It remains unclear whether more robust neural latent representations exist, particularly for implicit confidence. We address this by applying complementary machine learning approaches to comprehensive, brain-wide intracranial electroencephalography (iEEG) data acquired without explicit confidence reports. Using explainable feature selection methods, we elucidated key spatiotemporal iEEG components, especially in the frontal cortex, that readily distinguished confidence characteristics related to decision correctness, difficulty, and speed, and that were transferable across motor modalities. Despite inconsistencies in the features identified across methods and conditions, dimensionality reductions revealed robust low-dimensional neural latent representations. Furthermore, their neural trajectories within a shared subspace enhanced the distinguishability of confidence characteristics. Overall, our results uncover low-dimensional spatial and temporal neural latent representations underlying human implicit perceptual decision confidence that are robust to analytical methods and decision conditions.

## Introduction

Perceptual decisions are often made under uncertainty, and hence, are accompanied by a sense of confidence in estimating the correctness of a sensory informed decision [1–3]. Confidence, a form of metacognition, systematically covaries with fundamental properties of perceptual decisions, particularly sensory evidence strength, choice accuracy and reaction time [4, 5]. Such relationships imply that confidence need not be accessed exclusively through explicit subjective reports, an often used approach [6], which may interfere with decision process [7, 8].

Neural signature of decision confidence can offer implicit information. Electrophysiological recordings in behaving animals have shown that neuronal activity involved in forming perceptual decisions can simultaneously carry information about decision certainty [9–11]. Non-invasive neuroimaging and electrophysiological studies in humans have nevertheless implicated a substantially broader set of regions and signals. In particular, neural correlates of decision confidence have been observed across prefrontal, medial frontal and cingulate, insular, parietal and frontoparietal regions, as well as in centroparietal and error-related electrophysiological signals [12–18].

Indeed, intracranial EEG (iEEG) has recently revealed widely distributed perceptual decision signals whose amplitudes covary with sensory evidence, accuracy and reaction time [19–22], which can also be abstract, effector-independent representations [20, 21]. Yet, so far, iEEG study of implicit decision confidence is more hypothesis-driven with a focus on specific brain regions [23, 24]. Decision confidence may therefore not be localised to a single brain region, but instead emerge from distributed computations from apparently heterogeneous activities. However, the precise brain-wide neural organisation of confidence remains unresolved.

A complementary possibility is that robust confidence representations emerge not at the level of brain regions, but in the geometry of population activity. Neural population analyses and theories increasingly show that high-dimensional activity from heterogeneous activities can be organised along a small number of latent dimensions encoding behaviourally relevant variables and that can separate sensory, cognitive and motor variables into distinct directions or subspaces for convenient readout [25]. Whether implicit perceptual decision confidence is represented in such a low-dimensional organisation from brain-wide data remains unknown.

Here, we test these possibilities using comprehensive brain-wide human iEEG data combined with complementary explainable and dimension-reduction machine-learning (ML) approaches (Fig. 1A). Rather than requiring explicit confidence reports, we probe implicit confidence through three established characteristics of perceptual decision confidence, i.e. decision correctness, difficulty and speed, using data from a perceptual decision task with different effectors for choice reporting (Figs. 1B-C).

**Fig. 1.**
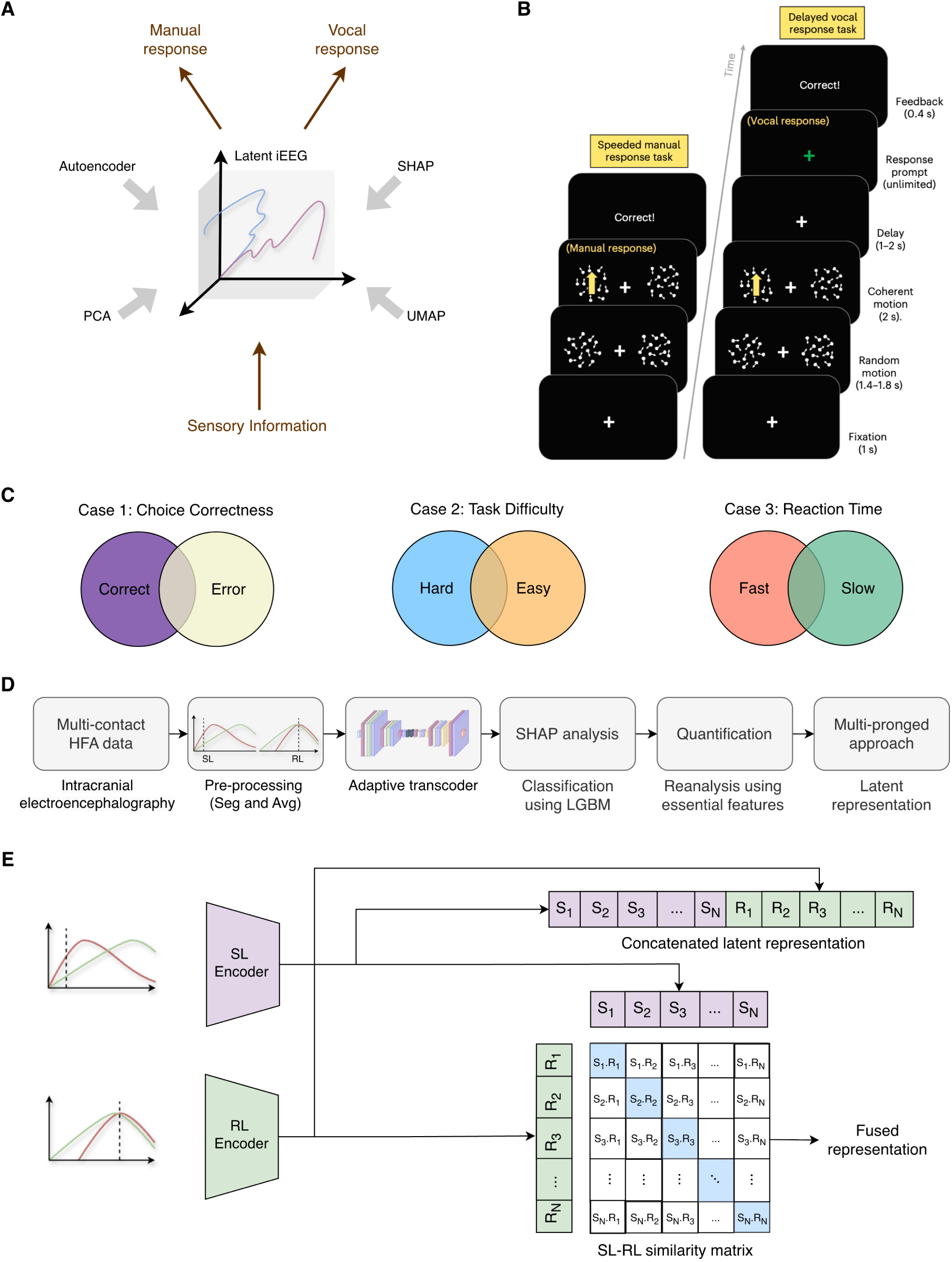
Data-driven approach to identify iEEG features and latent representations of perceptual decisions. **A.** Schematic of multi-pronged approach in elucidating latent iEEG representations, applied to different response modalities. **B.** Discriminating up-vs-down motion with different effectors. Adapted from [20]. **C.** Machine-learning (ML) classification on correct-vs-error choices, hard-vs-easy tasks (motion coherence levels) and fast-vs-slow responses (reaction times (RT) in manual response task). **D.** ML framework (see main text). Pre-processing include data segmentation (Seg) based on RT and trial-averaging (Avg) within fast or slow RT bins; done independently for stimulus- (SL) and response-locked (RL) data. **E.** Data fusion approaches from extracted SL and RL latent embedding (left): early SL-RL (ESR) fusion (right, top) and contrastive SL-RL (CSR) fusion (right, bottom).

We first evaluate whether time-compressed iEEG activity could predict these characteristics and identify the recording sites contributing most strongly to their classifications while agnostic to effectors. We then ask whether their spatially distributed neural signatures form a lower-dimensional representation that is more robust than electrode-level features across analytical methods and conditions. Finally, we test whether neural trajectories in latent space can enhance the separability of confidence characteristics (Fig. 1A). Various ML techniques were evaluated, including for robustness analysis (Figs. 1D-E). Together, we found that, despite heterogeneous and inconsistent activity across brain regions and analytical approaches, implicit perceptual confidence is organised within a predominantly common low-dimensional neural latent space whose trajectories provide more robust representations of decision confidence.

## Results

The iEEG data from this study was chosen based on its comprehensive coverage across brain regions in perceptual decision-making tasks with different effectors for choice reports [20]. Human participants in the study had to visually discriminate coherent up-vs-down motion direction, and they had to make speeded finger-pressing (manual) response (Fig. 1B, left), or delayed vocal response upon cue (Fig. 1B, right) to report their decisions. Further details can be found in Methods and the original paper [20].

Out of a total of 3, 380 iEEG contacts used, a subset was previously found to exhibit decision signals distinguished from those related more to sensory processing or motor execution (effector) - 233 effector-based contacts in manual response task, of which 84 contacts were from vocal response task, while only 2 sensory encoding contacts [20]. In comparison, 182 effector-independent decision-based iEEG contacts were identified by their involvement in evidence accumulation (ramping) neural dynamics independent of motor actions and sensory locations in manual response task, of which 111 contacts were from vocal response task [20].

We first considered the manual response version of the task with participants’ reaction time (RT) (Fig. 1B, left), as neural activity encoding evidence accumulation is known to be more closely aligned with the speed of free response [20]; the vocal response version of the task had no participants’ RTs. We began with the iEEG’s high-frequency activity (HFA) data in the manual task and preprocessed through segmentation of data based on RT (histogram-based binning) and averaging over trials, participants and cases in both stimulus-locked (SL) and response-locked (RL) neural activities, respectively (Methods) (Fig. 1D, left). The three cases for classification were: (i) Case 1 - correct-vs-error choices; (ii) Case 2 -hard-vs-easy task based on motion coherence difficulty level; and (iii) Case 3 - fast-vs-slow decisions based on individual RTs (Fig. 1C).

We initially directly applied principal component analysis (PCA) to the preprocessed neural data. For both stimulus-(SL) and response-locked (RL) activities, we found that the classes within each of the three decision confidence characteristic of decision correctness, RT and difficulty were inseparable in PCA space (Extended Data Fig. 1). Hence, more systematic and rigorous intermediate steps are needed. We then applied a suite of ML based selection (Figs. 1D-E) of iEEG features to distinguish classes within each confidence characteristic case (Fig. 1C) and identify the key temporal and spatial iEEG components. Further, a multi-pronged approach involving topological, nonlinear and linear based dimensional reductions were also employed (Fig. 1A).

### Temporally compressed SHAP-informed representations predict decision correctness, difficulty and speed

Using our previously developed temporal convolutional autoencoder (TCA) [26], we extracted temporal latent representations of the HFA data (Fig. 1D, middle), which compressed the original iEEG data in the temporal dimension to elucidate key temporal components while enhancing reconstruction capability (Supplementary Table 1). To reduce computational cost, we employed unsupervised adaptive transfer autoencoder, i.e. adaptive transcoder (ATC), which leverages pre-trained weights [27], on the TCA. ATC employs a type of transfer learning in which we used the network’s parameters (weights) trained on one case (e.g., Case 1) as the initial values for learning other cases (e.g., Cases 2 and 3). ATC attained optimal performance within half the training time as compared to the original TCA, without sacrificing reconstruction loss via various metrics. Hence, ATC was used for all subsequent analyses. ATC yielded a total of 64 time samples from the initial 256 in the neural latent (temporal) representation for each HFA sample.

Due to the inherent sparse and heterogeneous nature of the iEEG data [20, 22], with different participants having different contact coverages, we divided the data based on iEEG contacts (spatial features) instead of trials when conducting ML cross validations (CVs). Such a partial feature exclusion approach allowed assessment of whether the trained ML model could detect task-relevant differences while generalising to unseen combinations of contact features.

After temporal data compression, we sought to elucidate the most important iEEG contacts for (mis)classifications. To achieve that, we used an explainable ML method SHapley Additive exPlanations (SHAP) [28] (Methods) to remove spatial feature redundancies with low SHAP’s Shapley values (i.e. low contributions to classifications). When conducting SHAP, we evaluated it using three different CV strategies. First, we used leave-one-subject-out (LOSO) CV to obtain each participant’s individual SHAP rankings and then used their cumulative ranking (just acquiring 2% from each) to reevaluate them using stratified group 5-fold (SGKF) for post-hoc validation to discover the optimal contacts for good classification performance (see Methods).

Fig. 2A illustrates sample original (grey) and reconstructed trial-averaged HFA time-courses in specific brain regions and participants in each subcase. The brain regions shown were essential features, highly ranked and consistently identified across cases using SHAP’s Shapley value. The compressed signals could faithfully reproduce the original signals. Moreover, using gradient-weighted class activation mapping (Grad-CAM) [29], we found the ramping and post-response epochs (Figs. 2A, magenta) to be important, consistent with both the pre- and post-decision confidence hypotheses.

**Fig. 2.**
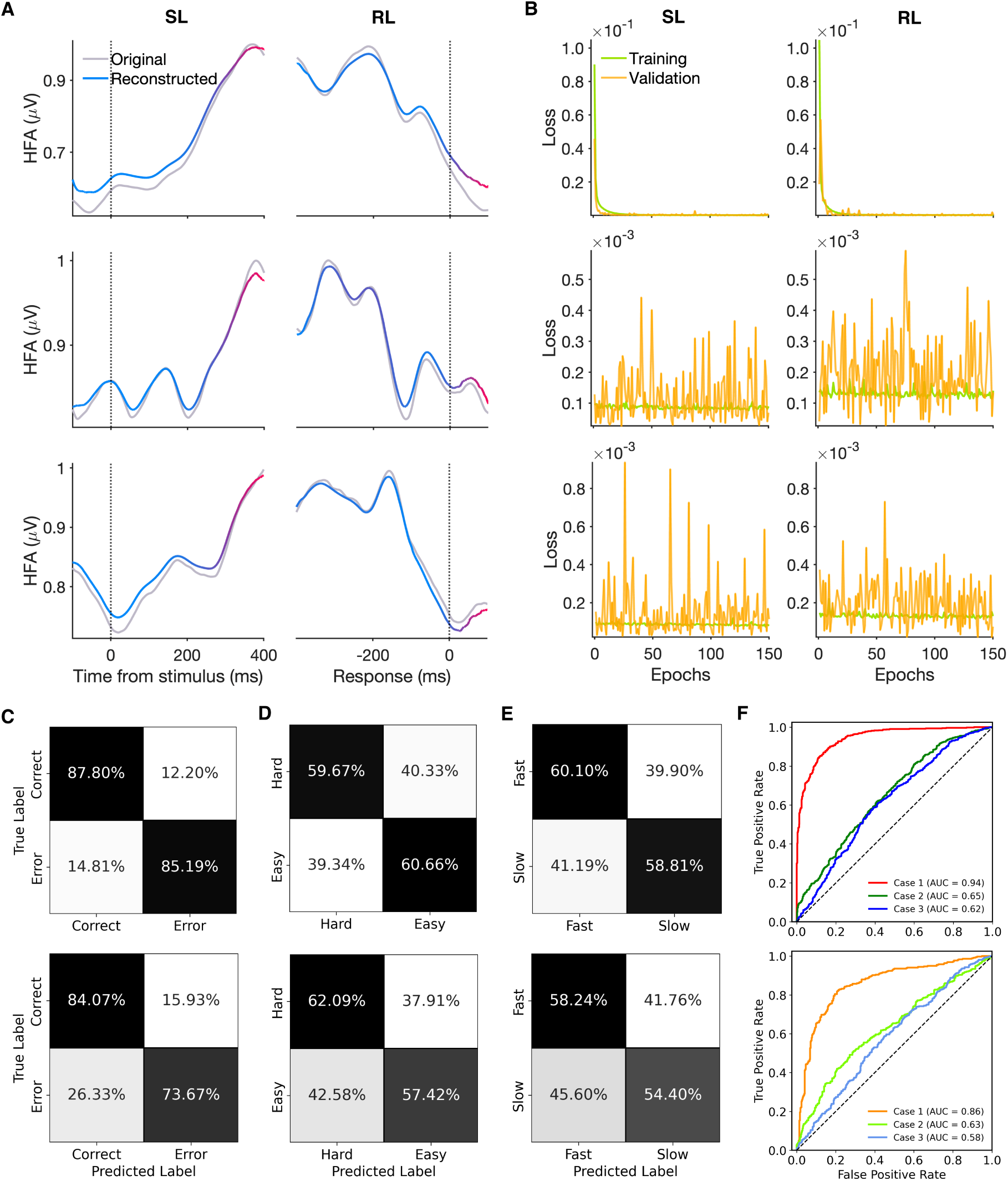
Stimulus- (SL) and response-locked (RL) HFA timecourses across (sub)cases and classification performance for SHAP-based feature selection after time compression. **A.** Sample SL and RL activities of individual participants S14 (top), S15 (middle) and S16 (bottom) for Case 1 (hard task subcondition) (top), Case 2 (slow RT subcondition) (middle) and Case 3 (hard task subcondition) (bottom), in manual response tasks. Original (grey) and corresponding latent representation reconstructed (normalised Grad-CAM overlaid) HFA timecourses, mapped from blue (low importance) to magenta (high importance). iEEG contact regions (top to bottom): right lateral occipital (S14-ROl8), right caudal anterior cingulate (S15-RFm4) and left caudal middle frontal (S16-LFp6) cortices. **B.** Loss plot across epochs of training (green) and validation (amber) of the ATC for Case 1 (top), Case 2 (middle) and Case 3 (bottom) for SL (left) and RL (right) HFAs, respectively. **C-F.** LGBM classification performances using 2% of ML selected features (top) and identified effector-independent features from the previous study [20] (bottom). Only for SL analysis on Cases 1 (C), 2 (D) and 3 (E). ROC curves and corresponding AUCs for different cases; based on features from 2% of SHAP-based (top) and original study (bottom) (F).

We observed very low loss attained during ATC training (green) and validation (amber) phases for all the cases (Figs. 2B). This offers assurance in classifying using different ensemble ML (EML) models (Methods). By aggregating across all iEEG contacts, our initial exploration found that the light gradient-boosting machine (LGBM) model [30] showed slightly better classification performance than the other EML models considered, while remaining comparatively less complex (Methods and Supplementary Table 2). Moreover, considering all iEEG contacts, we found TCA-based classification to attain higher accuracies than the original uncompressed version (Supplementary Table 3), justifying the earlier temporal compression method.

Critically, we found that 2% of the essential SHAP-based iEEG contact features could classify as well those of the 5%, 10% and the full data (Supplementary Table 4). We shall henceforth focus on using these 2% of the iEEG contact features, in their temporally compressed form. We will begin by performing ML classifications on the SL data. For Case 1, we found that the trained LGBM algorithm could classify correct from error choices with high accuracies (Fig. 2C, top). Although substantially above chance level, the classification performances on task difficulty (Case 2) and RT (Case 3) were lower than Case 1, while Case 3 performed the worst (Figs. 2D-F, top). This might reflect that the identified contacts were more internally generated with stronger affiliation to correctness than sensory or motor processing.

To validate this, we used the 182 effector-independent contacts identified in the previous study [20] and found a similar pattern across the cases (Figs. 2C-F, bottom). Here, we followed the same processing pipeline but without applying any ML-based feature selection processes. Additionally, these selected contacts generally had classification performance comparable to the SHAP-identified ones (Figs. 2C-F, compare bottom against top).

In addition to the analysis on SL data, we also analysed RL and two types of SL-RL fused data. The latter comprised a näıve early SL-RL (ESR) fusion baseline, and an advanced contrastive SL-RL (CSR) fusion (Fig. 1E) (Methods). For Case 1, significant differences were observed in SL, RL and ESR (*p <* 0.001 in most metrics; Cohen’s *d* ranging from 1.04 to 1.69), whereas for Case 3, stronger effects were observed particularly in RL, ESR and CSR (*p <* 0.01 in most metrics; Cohen’s *d* ranging from 0.71 to 2.06). For Case 3, our approach achieved higher classification performance than that with features from previous study [20]. However, for Case 2, our ML-based approach showed comparatively smaller differences overall, with generally low effect sizes in SL and mixed results across RL, ESR and CSR. Hence, there is variability across methods. For additional metric evaluations, see Supplementary Tables 5 and 6, which used: (i) all available HFA features, (ii) 2% essential SHAP features acquired from ATC, and (iii) features identified by Gherman et al. [20].

### Distributed SHAP-identified brain regions can distinguish decision confidence characteristics and are effector-independent

We next aim to reveal the specific brain regions of the top 2% SHAP-ranked iEEG contacts, their relative contributions to classifications and their consistencies across cases and analyses. We first inspected their spatial overlaps. For Case 1, the most important brain regions, ranked from the highest, were in the frontal, parietal and temporal cortical regions in SL and CSR based analyses. This order was slightly different for the RL and ESR analyses, in the order of frontal, temporal and parietal cortical regions. Similarly, in Cases 2 and 3, the features were across brain regions in all the analyses, based on ranking. Hence, they are distributed across the cortex, with the frontal region showing the most consistent contribution.

Upon closer inspection, we found a similar regional pattern in Case 1, with iEEG contact contributions distributed primarily across frontal regions in SL (43.44%), RL (36.14%), ESR (42.28%) and CSR (44.31%) analyses, followed by parietal in SL (22.81%) and CSR (28.62%) and temporal in RL (30.22%) and ESR (24.07%) analyses (Fig. 3A). Comparatively, there were smaller contributions from the insula and occipital regions. See Fig. 3A for other contributing regions. Cases 2 and 3 were also mainly distributed across the frontal, parietal and temporal regions across analyses, but showed a more consistent contact-level pattern than Case 1 (see Supplementary Note 1.1).

**Fig. 3.**
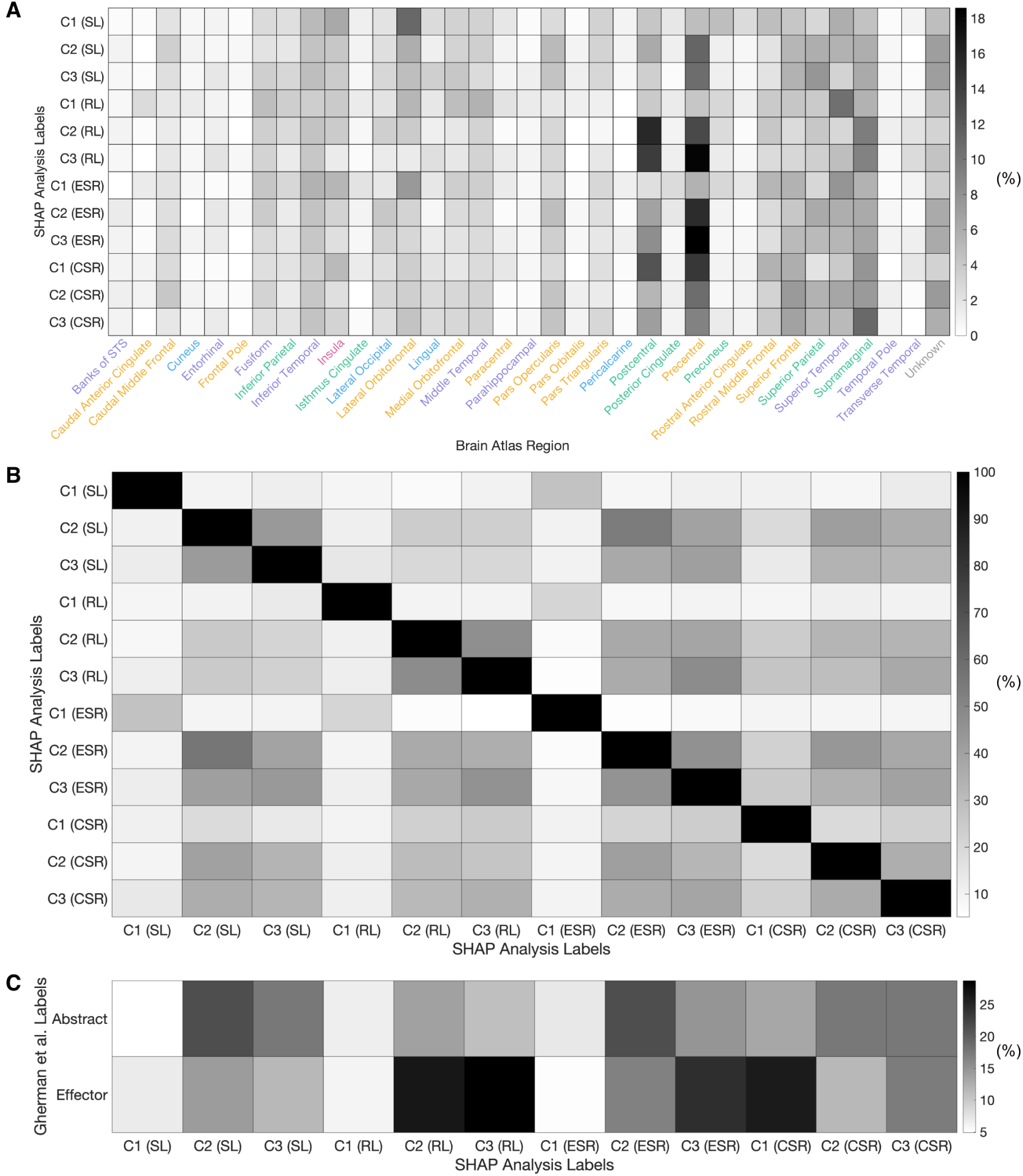
SHAP-identified essential iEEG contact locations. C1, C2, and C3: Cases 1, 2, and 3, respectively. **A.** Brain region distribution of individual analyses. Colours: frontal (amber), insula (magenta), occipital (blue), parietal (mint), and temporal (violet) brain lobes. **B.** Comparison (% overlap) of SHAP analysis (essential 2% features). Matrix is asymmetric as each entry shows the fraction of the row feature list that also appears in the column list. **C.** Comparison (% overlap) between previous study [20] and SHAP analysis (essential 2% features). Effector-independent and effector-dependent contacts denoted by Effector and Abstract (abstracted from effector), respectively.

At the individual contact locations, we found overlaps in Case 1 to be limited, suggesting a more distributed and inconsistent spatial pattern (Fig. 3B). In contrast, Cases 2 and 3 showed greater overlaps with each other across analyses, with pairwise overlap values of 42.31-43.28% in SL, 46.74–47.89% in RL, 45.45–46.08% in ESR and 35.35–36.27% in CSR (Fig. 3B). The iEEG contacts identified from the intersection among the three cases were mainly distributed across the frontal and temporal regions, with the frontal region showing the largest overlap in most analyses (Supplementary Note 1.1).

At the subregion level, the contacts shared across the three cases were relatively sparse and differed across different analyses (Supplementary Note 1.2). The largest overlap was observed between RL and CSR (22.22% from RL to CSR and 11.11% from CSR to RL), whereas SL showed only small overlap with RL and CSR (5.56–16.67%) and ESR showed no overlap with CSR (Fig. 3B). This suggests that although a few common contacts were preserved across subcases, the specific contact-level representation remained strongly dependent on the analytical setting. The inconsistency might be due to the inherent sparse and heterogeneous iEEG recordings, or pointing towards deeper and more robust neural representations yet to be found.

Next, for each case, we search for any intersection of the identified iEEG contacts with those identified in the previous study [20]. We found our top SHAP-ranked features to partially overlap with the labels reported in the previous study [20]. Across the analyses, the overlaps with their effector-independent labels ranged from 4.9% to 21.4%, whereas the overlaps with their effector-dependent labels ranged from 5.2% to 28.8% (Fig. 3C) (See Supplementary Note 1.3). The limited overlaps while still having similar classification performances (Figs. 2C-F) may indicate that these selected contacts coordinate to form part of some underlying computation.

We now sought to validate, via the SL data, whether the ML model trained on one effector type (e.g. manual response) could generalise to another effector type (vocal response) in a zero-shot setting (ZL), i.e. without further training. Since there was no actual RT data in the vocal response task, we restricted this comparison to Cases 1 and 2 and SL data. For ZL, the model generalised manual-to-vocal responses better than vocal-to-manual responses. Specifically, for Case 1, the manual-to-vocal setting achieved 0.871 *±* 0.013 accuracy, 0.869 *±* 0.009 F1-score and 0.953 *±* 0.003 AUC, whereas the vocal-to-manual setting yielded 0.863 *±* 0.004 accuracy, 0.861*±*0.007 F1-score and 0.936*±*0.004 AUC. A similar trend was observed for Case 2, albeit lower accuracies (*∼* 0.53-0.56), where manual-to-vocal performance remained marginally (*∼* 2%) higher than vocal-to-manual performance across all three metrics.

Overall, the trained model in the manual response task could directly be used for classification in the vocal response task without additional training. This suggests that the top 2% SHAP-ranked iEEG features were effector independent, i.e. the learned feature representation retained discriminative information that could be directly transferred across the two choice reporting modalities. However, the highly distributed neural signals identified and the lack of consistency might indicate additional features beyond those found at the brain regional level.

### Robust low-dimensional neural latent representations across decision confidence characteristics

Building on the latent temporal representation and the top 2% SHAP-ranked iEEG data, we sought to further elucidate its latent representations. First, we used Uniform Manifold Approximation and Projection (UMAP) [31] for nonlinear, topological based dimensional reduction (Methods). For SL data, we found that the latent space captured discriminative features for each case and were embedded within much lower dimensions than the original HFA spatial dimensions (Figs. 4A-C). It should be noted that each time-compressed activity time points were mapped as individual data points in UMAP space. The distinctive clusters in latent space (Figs. 4A-C) in distinguishing the classes in each case lends support to the good classification performances of the considered cases (Figs. 2C-F). Other analyses with UMAP gave similar results (Extended Data Fig. 2), hence demonstrating robustness.

**Fig. 4.**
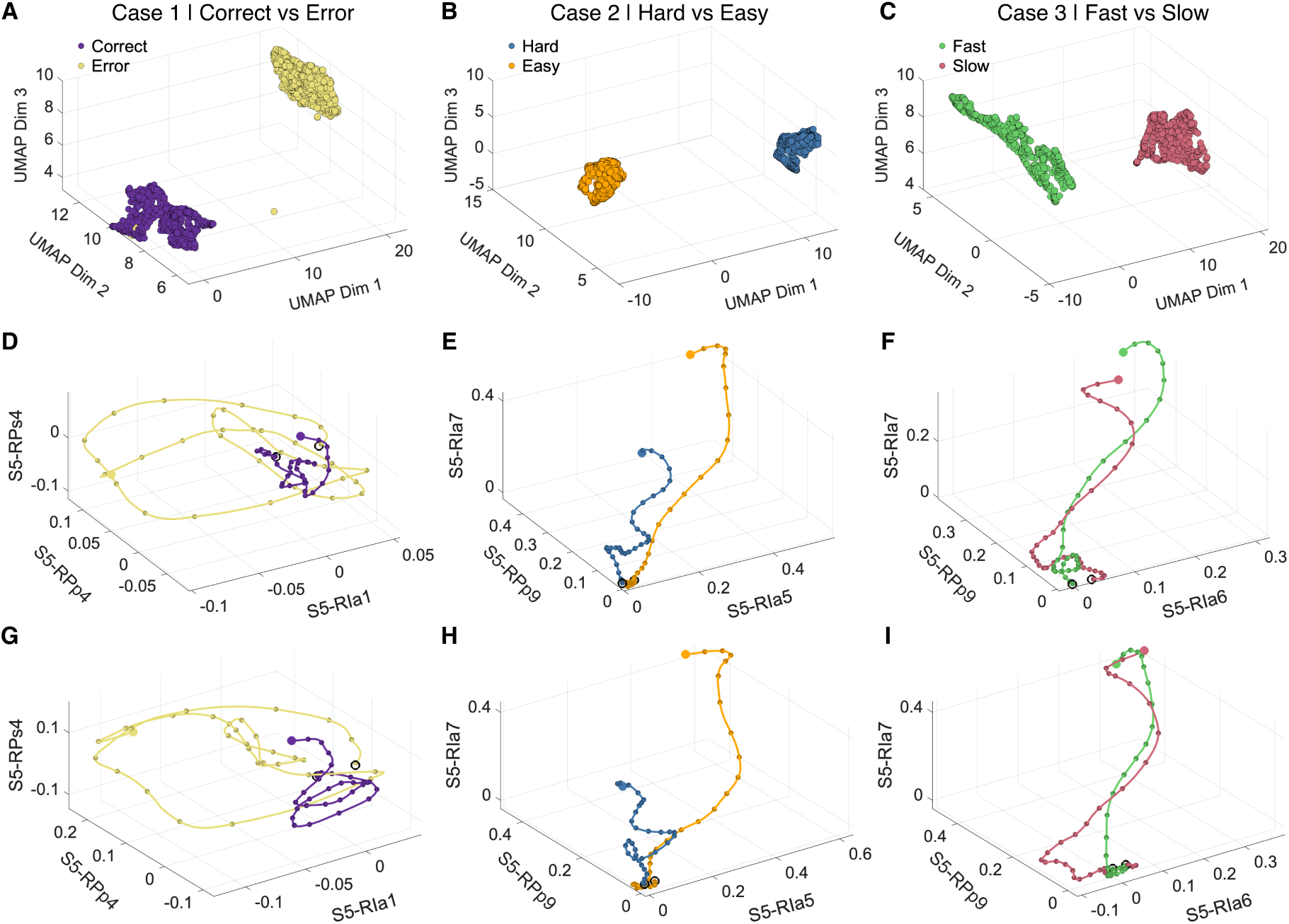
Low-dimensional latent representations from ATC of iEEG HFA in SL manual response task. Left, middle and right panels: Cases 1, 2 and 3, respectively. **A-C.** 3D visualisations of UMAP projection for the three cases. **D-F.** Trial-averaged neural trajectories projected onto SHAP-based high-ranked iEEG contacts as feature dimensions for all the cases for sample participant 5. **G-I.** Same trial-averaged neural trajectories with additional sub-conditions of easy task (G and I), and slow RTs (H). D-I. Open circles: starting points; filled circles: end points. Colour labels the same as A-C.

To determine whether simpler dimensional projection method offers more explainable and meaningful representations, we project trial-averaged neural trajectories of the original HFAs onto the spatial feature, i.e., contact location, space spanned by three high-ranked SHAP-based iEEG contacts for each case. Figs. 4D-F demonstrated samples of clearly distinguishable trajectories within each case for a single participant (participant 5), consistent with the observed high classification accuracies. Interestingly, the trajectories in Cases 1 and 2 looked more separable than that of Case 3, consistent with the classification trends in Fig. 2C-F. A similar pattern was observed for the sub(cases) (Figs. 4G-I), indicating the robustness of these trajectories regardless of specific sub-conditions (Extended Data Fig. 3 shows the remaining sub-conditions). However, these results were based on separate (sub)cases.

Fig. 5A illustrates the brain topographical map, co-registered with individual participant’s structural MRI data, for the above SHAP-identified essential contacts for SL (top) and RL (bottom) analyses in manual response task (Supplementary Tables 7 and 8 shows detailed contact locations). This again showed the diversity of the brain regions involved in each case. Yet, there were overlaps (18 iEEG contacts) across all the considered cases (grey regions), indicating these locations of intersection may possibly encode neural signals not confined to decision correctness, difficulty or speed.

**Fig. 5.**
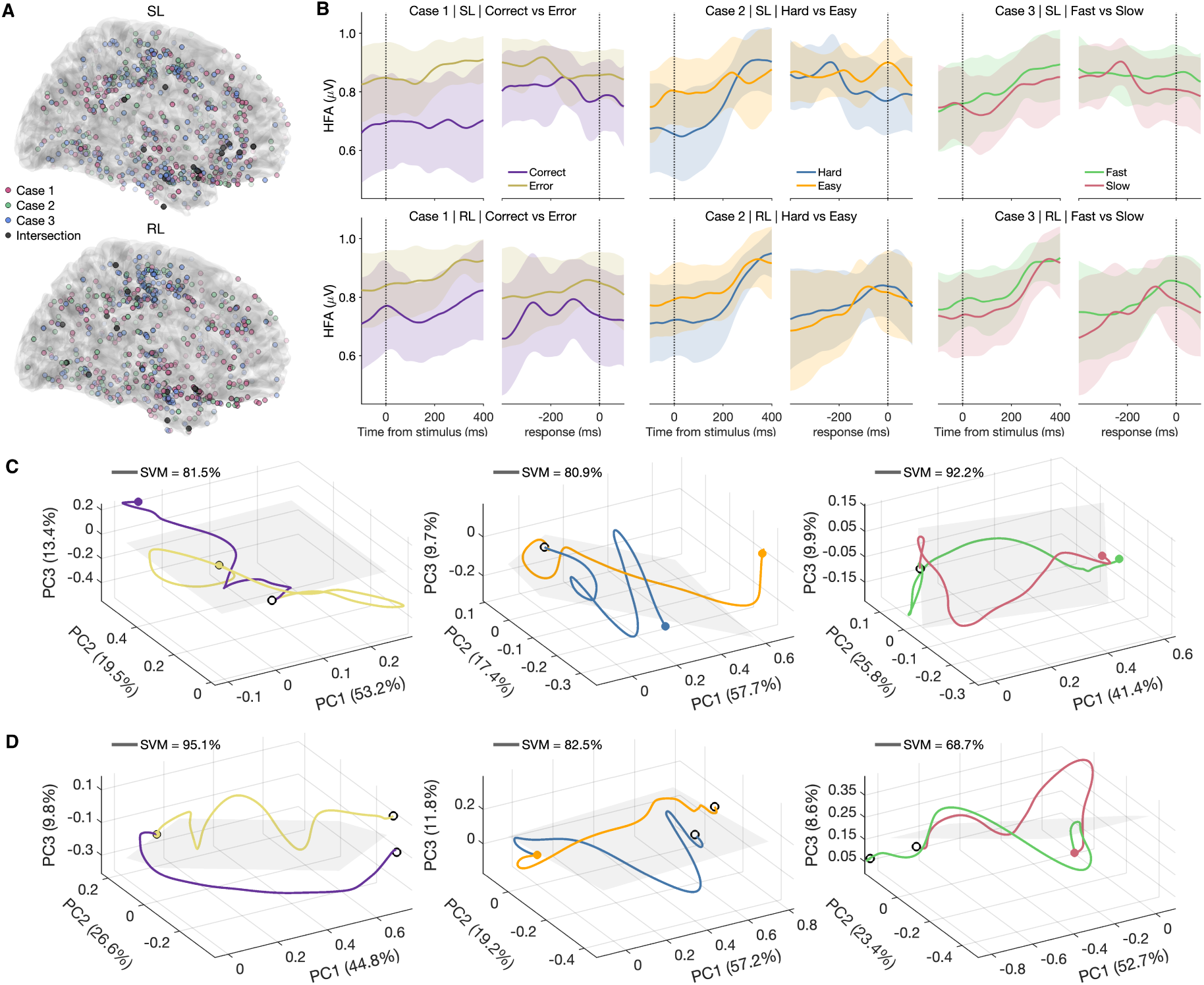
SHAP-identified contact locations and grand-averaged activities for SL and RL analyses. **A.** Dark grey: Projected structural MRI topographical plot mapping essential HFA contacts (Case 1 (red) *∩* Case 2 (green) *∩* Case 3 (blue)) identified by SHAP in the SL (top) and RL (bottom) analyses. Grey: 18 (18) contacts overlap across all cases for SL (RL). Colour transparency encodes SHAP importance (more opaque denotes higher importance). **B.** PCA-weighted grand-averaged neural activity timecourses; SL (top) and RL (bottom) analyses. **C–D.** Grand-averaged neural trajectories projected onto PCA space of the SHAP-identified top-ranked features that intersected (A, grey regions) across all three cases. **C,** SL analysis; **D,** RL analysis. Left to right: Cases 1, 2 and 3, respectively. Linear SVM decision hyperplanes (grey planes) overlaid for further classification of neural trajectories. Open circles: starting points; filled circles: end points. Colour labels the same as in B.

Fig. 5B showed sample cortical contact SL (top) and RL (bottom) PCA-weighted grand-averaged activity timecourses for the different cases. In general, for SL activity, we could observe stereotypical neural activities that ramped up over time, consistent with evidence accumulation hypothesis. However, when locked to their respective response time window [*−*400 to 100 ms] (RL), these activities did not exhibit clear modulation by decision correctness, difficulty or speed, unlike in previous iEEG studies [20, 21]. There were even less modulatory effects when we performed näıve averaging of activities (Extended Data Fig. 4). This points towards possible coordinated computation.

Thus, we performed PCA and projected the grand-averaged SL activities of the intersected contacts onto PCA spaces. Strikingly, we found that about 80% of the data variance could be accounted for by the first three principal components (PC1, PC2 and PC3) (Fig. 5C). When we performed linear support vector machine (SVM) classifications of the neural trajectories in PCA spaces, we could classify the classes in each case with very high accuracies, with 81.5% for Case 1, 80.9% for Case 2 and 92.2% for Case 3. Similarly, for the RL analysis, SVM classification accuracies were 95.1%, 82.5%, and 68.7% for Cases 1-3, respectively (Fig. 5D).

We repeated with ESR and CSR analyses (Supplementary Notes 2.1, Extended Data Fig. 5A) and found that the number of contacts that overlapped across all cases to vary: there were 10 for ESR and 36 for CSR (Supplementary Tables 9 and 10 for details of contact locations). Interestingly, although these data fusion approaches exhibited clearer ramp-to-threshold-like patterns near response time than SL or RL analyses (Extended Data Fig. 5B), they had lower data variance accounted in PC1-PC3 and classification accuracies (Extended Data Figs. 5C-D).

As PC1 accounted for about half the data variance, its loadings were computed to identify key iEEG contacts’ contributions. We found that with SL or RL analyses, all three cases generally had their top-ranked iEEG contacts overlapping, ignoring the flipping of their loading signs (Figs. 6A-B). This indicates the neural latent subspace encoding decision correctness, difficulty and speed shared the same subspace as supported by the PC1 unit vectors projected onto the top three iEEG contacts for the three cases (Figs. 6C). With ESR and CSR, the loadings were relatively less similar across cases, explaining their lower classification accuracies. Further, CSR involved many iEEG contacts, and hence required more dimensions to capture data variance and the neural trajectories were noisier (Extended Data Fig. 6).

**Fig. 6.**
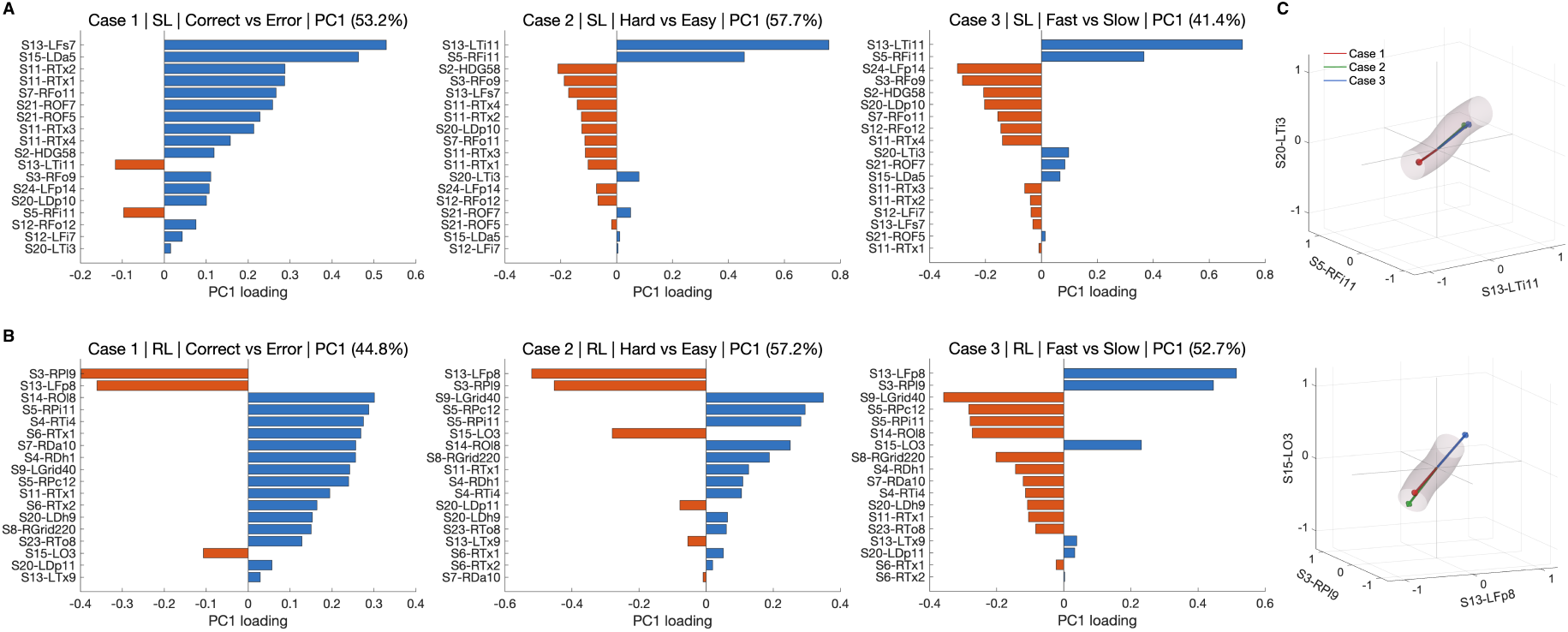
PC1 loadings reveal shared contact rankings across decision characteristics. PC1 loadings of essential iEEG contacts for stimulus-locked (SL) (**A**) and response-locked (RL) (**B**) analyses, computed from SL and RL data, respectively. **C.** PC1 loadings of top-three representative contacts for Cases 1–3 in SL (top) and RL (bottom). Grey tubes denote aligning along common manifolds.

We next quantify the neural trajectory characteristics in PCA by computing their path lengths, average speeds and displacements across all four analyses (Fig. 7). Here, path length represents the total distance travelled through PCA space, average speed represents the mean distance travelled per trajectory step, and displacement represents the Euclidean distance between the initial and final points of the trajectory.

**Fig. 7.**
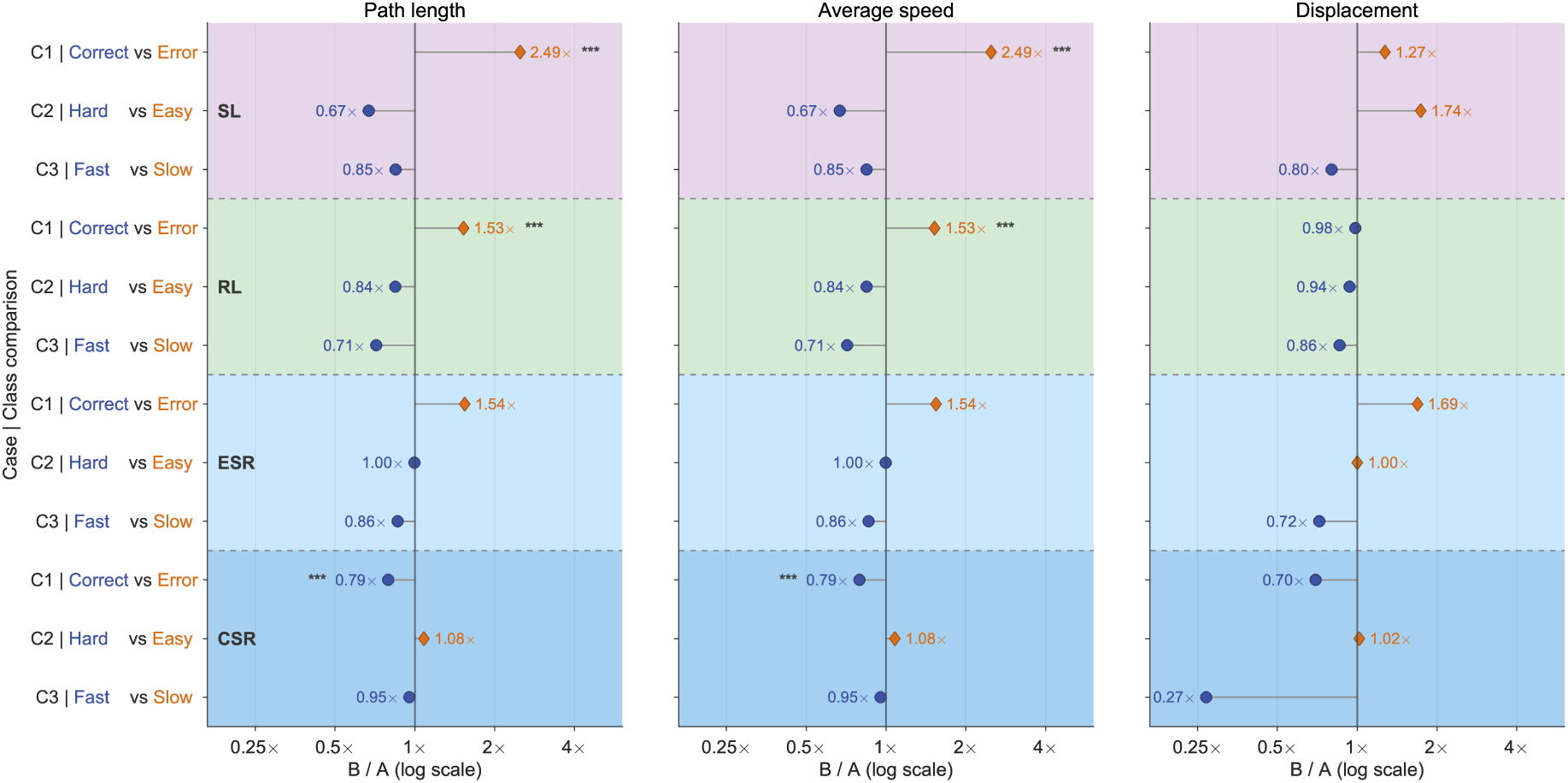
Characteristics of grand-averaged PCA neural trajectories across analyses. Left to right: path length, average speed and displacement, respectively. Top to bottom: stimulus-locked (SL), response-locked (RL), early SL-RL (ESR) fusion and contrastive SL-RL (CSR) fusion, respectively. Circles and diamonds indicate the two classes within each case, with horizontal lines joining each comparison. Marker values calculated directly from grand-averaged trajectories, whereas statistical significance was assessed across paired contacts using two-sided sign-flip permutation tests. Asterisks: Holm-adjusted significance.

Error neural trajectories’ path lengths were found to be significantly longer than correct trajectories in SL (2.49*×*) and RL (2.49*×*), and slightly shorter in CSR (0.79*×*) analyses (all *p*_adj_ *<* 0.001), but not in ESR analysis (Fig. 7). Average speeds followed a similar pattern to path lengths. For displacement, no class-level comparison reached significance. (Subclass-level trajectory metrics are illustrated in Extended Data Figs. 7 and 8). Thus, path length and average speed of neural latent trajectories could robustly distinguish correct from error decisions in SL, RL and CSR, although the direction of the effect differed in CSR (Fig. 6C).

## Discussion

Despite extensive research on perceptual decision confidence, the field has largely been dominated by perspectives at the neuronal, local neuronal population or brain regional levels [2, 3, 9]. However, the search for its neural correlates has yielded variable and sometimes inconsistent findings. Inspired by recent neural latent state studies in animal perceptual decision-making [25, 32], we proceeded to search for such an underlying neural state within comprehensive, brain-wide iEEG data from a perceptual decision-making experiment conducted without overt subjective confidence reporting. Rather unexpectedly, we uncovered predominantly very low-dimensional neural latent representations that could accurately and robustly distinguish human decision correctness, difficulty and speed, hence encoding key characteristics of implicit decision confidence.

These results were achieved using a complementary suite of ML techniques on heterogeneous and fragmented iEEG data [20]. First, our previously developed TCA compressed the iEEG activity time points, while explainable ML Grad-CAM method identified key time points that contributed the most to activity reconstruction. We found pre-decision ramping activity and post-decision activity to contribute the most (Fig. 2A), consistent with the evidence accumulator hypothesis of decision confidence while also pointing towards the importance of both pre- and post-decision times in decision confidence processing (e.g. [15, 33]).

Second, we applied explainable ML SHAP method, in combination with LOSO CV to account for inter-subject heterogeneity and sparse sampling, on top of the time-compressed activity to identify key spatial features, i.e. iEEG contact locations. We found that the top 2% of the contacts sufficed to readily classify the three decision confidence characteristics of decision correctness, difficulty and speed (Figs. 2C-F). Interestingly, distinguishing decision correctness was the most accurate. As the sense of correctness is a stronger form of metacognition [2, 4], and decision speed reflects internal decision commitment rate [32], while difficulty is driven more by external stimulus (evidence strength), this may indicate that the identified contacts could be more involved in higher metacognitive processing. Further supporting this proposition, we used zero-shot transfer learning, i.e. without additional learning and found that these reduced spatiotemporal iEEG features were generalisable across two different motor modalities (finger pressing and vocal response) in decision reporting, i.e. agnostic to effectors.

The contacts we identified were distributed across multiple brain regions (Fig. 3A). The more consistently identified contacts in the frontal cortex were consistent with the metacognition literature [2]. Moreover, the identified frontal and temporal regions were consistent with a previous iEEG study of implicit decision confidence [24]. However, spatial overlaps were limited across our different analytical methods and with those encoding effector-independent accumulator decision signals identified from previous study using the same data (Figs. 3B and C) [20]. These inconsistencies might be due to the nature of iEEG data collation, or more fundamentally, the distributed and varying nature of decision processing; perhaps the source of variability and sometimes inconsistent findings in decision confidence research [13–18]. Therefore, a more robust way to quantify neural measures of decision confidence is needed.

To address these issues, and encouraged by the low-dimensional neural trajectories projected onto top SHAP-ranked contacts and in UMAP (Fig. 4 and Extended Data Figs. 2 and 3), we performed linear PCA dimensional reduction, to elucidate a more robust and interpretable neural latent space. Remarkably, these neural representations consistently collapsed into low-dimensional manifolds (*∼*80% of data variance captured by the first three PCA dimensions) across analytical methods and conditions (Figs. 5C-D; Extended Data Figs. 5C-D). The neural trajectories were also qualitatively similar across (sub)cases (Extended Data Figs. 9 and 10). This should be compared with the absence of separability in grand-averaged activities (Figs. 5B and Extended Data Fig. 4), and in our earlier unsuccessful PCA attempt (Extended Data Fig. 1). Crucially, the neural latent subspaces for the three decision characteristics overlapped, hence sharing the same geometrical computation (Figs. 5A and 6).

Probing into the neural trajectories embedded within these low-dimensional latent spaces, we were able to discriminate the classes substantially more accurately using linear SVM method (Figs. 5C-D and Extended Data Fig. 5C-D). This lends further support that the lower dimensional representations are better neural representations. Moreover, by comparing three measures of neural latent trajectory, namely, its path length, average speed and displacement, we found latent path length and speed could predict decision correctness consistently across (SL, RL and CSR) analyses (Fig. 7). This should be compared to a previous iEEG study which showed that the latent activity timing of ventral visual and inferior frontal cortices were linked to decision time [24]. Yet, an animal study has shown that there is an internal decision commitment time that is not strongly linked to overt reaction time [32].

The findings in our study suggest that perceptual decision confidence may be governed by robust, low-dimensional neural latent dynamics, with heterogeneous cortical activity reflecting projections of this underlying neural organisation. This raises the possibility that distributed brain regions collectively coordinate to maintain stable decision-related dynamics even when source-level encoding appears inconsistent; this speculation deserves further investigation. Related human intracranial studies have also identified confidence-related signals but were more confined to specific brain regions of interest, particularly in the ventral visual cortex and pre-supplementary motor area (pre-decision confidence) and insula (post-decision confidence) [23, 24]. Compared to these studies, our work did not select specific brain regions *a priori*, or explicitly search for evidence accumulation signals, but was data-driven using machine-learning techniques.

Methodologically, this study introduces several advances. We jointly performed spatial and temporal feature selection, in contrast to most neuroscience approaches that focus on spatial features alone (e.g. [24, 25, 32]). Further, our cross-validation strategy operated over features rather than trials, better accommodating the sparse and heterogeneous nature of iEEG data. We employed SHAP for interpretable feature selection, bridging the gap between predictive performance and neuroscience insight. Additionally, we incorporated (ATC) transfer learning to enable efficient model training with a standard laptop, offering wider accessibility. Moreover, we introduced complementary temporal perspectives for data analyses. Rather than restricting the analysis to a single temporal alignment setting (SL or RL), we additionally employed data fusion strategies (ESR and CSR) to provide a more comprehensive perspective. Finally, our zero-shot cross-effector generalisation provides a stringent demonstration of effector-independent model robustness.

A limitation of our study is that there is no subjective confidence report or behaviour to validate the latent neural dynamics. This will be explored in future work. It will also be important to relate how these latent dynamics at source level is related to more established electrophysiological signatures at scalp level, such as the centroparietal positivity (CPP) [15, 34]. It should be noted that our study did not exclude the contribution of ancillary processes that may co-occur with decision processing, such as attention [35], motor preparation [36], or physiological artefacts [37]. Such contributions also likely reflect the inherently integrated nature of brain function. Further work, including novel task design, would be required to disassociate these factors.

In summary, this work complements hypothesis-driven approaches by leveraging data-driven methods to reveal robust low-dimensional neural dynamics underlying perceptual decision confidence. Our findings highlight the existence of abstract, effector-independent representations and suggest that distributed neural activity may reflect coordinated projections of a shared latent process. This framework opens new avenues for understanding human metacognitive computation and provides a foundation for future work at the intersection of NeuroAI and neurotechnology.

## Methods

### Data description

The dataset used in this study was based on a previous iEEG study [20], which involved epilepsy patients undergoing invasive electrophysiological monitoring. The experiment involved visual motion discrimination of the coherent direction of random-dot stimuli, and participants responded either with a speeded button press manually (*n* = 24), or vocally after a randomised delay (*n* = 12) (Fig. 1B).

The iEEG electrodes were implanted in participants as part of their clinical treatment for epilepsy. The exact placement was clinically determined, which provided extensive coverage of various brain regions but was not uniform across participants. Initially, 5636 intracranial sites were considered for analysis. After excluding 1572 contacts due to epileptiform activity or proximity to seizure zones, and a further 684 contacts located in white matter, a total of 3380 contacts remained for the final analysis. These included 2946 depth contacts, 273 grid contacts, and 161 strip contacts.

### Data pre-processing

The iEEG data, originally recorded at 1 kHz and 3 kHz, was resampled to 512 Hz and epoched from *−*3 to 3 seconds relative to sensory stimulus onset time. High-frequency activity (HFA) (70-170 Hz) was selected for consistency with previous iEEG work.

For the manual version of the task, the HFA data were segmented into fast and slow trials by binning reaction times (RTs), following previous work [38] to capture graded evidence accumulation neural activities. First, trials with RTs exceeding a set threshold (calculated as the number of samples divided by the sampling frequency) were removed to eliminate outliers. RTs were then divided into two bins using histogram-based binning: trials with RTs below the first bin edge classified as fast, and those above the second bin edge as slow. This segmentation allows us to compare neural activity across different response speeds to provides insight into decision speeds.

The length of the HFA data for each participant varied considerably in the manual task. Hence, we considered a fixed window of size [*−*100 to +400ms] with respect to stimulus onset for SL, and [*−*400 to +100ms] with respect to response time for RL, in order to generalise over different participants and (sub)cases.

Motion coherence levels of the stimulus were categorised as hard or easy based on participant-specific perceptual thresholds, determined before the experiment. The hard trials generally had lower coherence, making motion direction harder to discern, whereas easy trials had higher coherence. The low coherence level was set at half the high coherence level to maintain a meaningful contrast [20]. The HFA data were averaged across all trials individually for each participant and normalised to remove any inherent scaling differences (Fig. 1D, left).

### Explainable and interpretable machine learning

*Temporal convolutional autoencoder (TCA):* The TCA model is adapted from our previous study [26]. The TCA comprises an encoder and decoder network, which is trained to learn the compact hierarchical representations of the input HFA data (Fig. 1D, middle). Extended Data Fig. 11 depicts the detailed system architecture of the TCA model. See Supplementary Note 3.1 for the mathematical formulation behind the proposed TCA framework.

The objective of TCA is to find the optimal weight parameters to minimise the cost function. In our implementation, the Adam optimisation technique was used with a learning rate of 10*^−^*^3^. Leaky ReLU was used as the activation function, with a negative slope parameter (alpha) of 0.2, allowing gradient propagation through negative values.

To visualise the 64-dimensional compressed latent space representation, we used UMAP [31] to reduce the data to three dimensions. In our simulation, we set *n neighbors* = 15 (the number of nearby points used to determine local structure) and *min dist* = 0.1 (the minimum distance allowed between each point in the low-dimensional space). This configuration revealed distinctive clusters corresponding to the different classes, despite the TCA being trained only for reconstruction.

*Adaptive Transcoder (ATC):* To improve efficiency and reduce computational cost, we used an adaptive training strategy for the TCA. We started the model training with weights learned from previous training. This transfer learning approach led to the convergence of the loss with reduced training time and allowed the model to adapt more effectively to new datasets. The TCA was initially trained for 300 epochs for Case 1 and the weights were then used to retrain for 150 epochs for Case 2 and Case 3 , respectively. However, the number of trainable parameters remain the same for both models, with 38, 432 parameters in total, including 38, 296 trainable and 136 non-trainable parameters. Since classification was performed individually for each case, any potential issues like bias transfer or misalignment between datasets were avoided, ensuring both efficiency and reliability.

*Gradient-weighted Class Activation Mapping (Grad-CAM):* Grad-CAM is employed in the ATC pipeline to provide post-hoc visualisation of where the model focuses during SL and RL analyses. In this setting, Grad-CAM generates activation heatmaps using gradients with respect to the final convolutional features of the ATC, thereby highlighting the input time-series that contribute most strongly to the learned reconstruction-related representations [29]. Since this can be implemented without altering the ATC architecture, it serves as a seamless addition for explainability by providing intuitive visual information for assessing the temporal regions most relevant to the model’s internal representations.

*SHapley Additive exPlanations (SHAP):* SHAP provides a unified approach for interpreting ML models by quantifying each feature’s contribution to the model’s predictions [28]. It uses a game-theoretic framework to assign importance values to individual features, ensuring transparency and interpretability even in complex models. In our study, SHAP was applied to the LGBM classifier trained on the latent representation of HFA data to identify the most influential features in the classification process.

### Fusion approaches

Neural activity is often analysed with respect to external referents, such as the onset of a sensory stimulus (SL) or an overt motor action (RL). Building on our initial work [22], we redesign a pipeline so that it can view the same neural process from different complementary (SL and RL) angles simultaneously. Here, we first learn latent embeddings from SL and RL neural data separately, and fuse them together. Fusion introduces a new perspective by combining both latent embeddings into a single shared latent representation, allowing the downstream ML classification model to leverage information tied not only to stimulus onset, but also to the response.

Two fusion methods are employed: early fusion (ESR) and contrastive fusion (CSR). ESR is a strategy that combines multiple data sources or modalities right at the input level or initial processing stage before feeding them into a single model. In comparison, CSR combines multiple data types or views by pulling similar combined features close together and pushing different ones apart in a shared mathematical space. See Supplementary Notes 3.2 and 3.3 for details.

As an ablation, we trained the same fusion frameworks directly on the uncompressed time series (without using ATC). This experiment was motivated to test whether SL-RL alignment can be learned without an explicit time-compression step, since the input dimensionality and representation capacity differ substantially from the ATC-based embedding pipeline. From the results obtained (Supplementary Table 3), it is notable that the latent embeddings learned by ATC help the model learn and fuse the information more intuitively, resulting in better discrimination performance.

### Model classification and performance

After obtaining the latent space representations from the ATC model, we used these features as inputs to a series of ensemble ML (EML) classifiers to evaluate their discriminative performance across all cases. To ensure independence and eliminate potential biases, the dataset was first annotated and shuffled. The encoder output of the trained ATC was used as input for various EML classifiers, comprising random forest classifier, adaptive boosting, light gradient boosting machine (LGBM), eXtreme gradient boosting, and soft voting ensemble classifier. The data were split into training (68%), validation (12%) and testing (20%) sets. The classifiers were trained using the training and validation data and evaluated on the testing data with stratified 10-fold (SKF) CV initially. Among all the classifiers, LGBM consistently delivered the best performance and was selected for further analysis.

To make the ML pipeline more robust, we evaluated three different CV approaches (Supplementary Table 3) for ML model training and validation. This was because, during our initial analysis a few participants disproportionately influenced the feature rankings across folds, leading to partial comparisons. This was later overcome by employing LOSO CV, in which we selected only the essential top 2%, 5% and 10% features and performed reclassification with SGKF CV. The main intuition behind this was to determine the smallest subsample of high-dimensional data that would still yield decent classification performance. We re-analysed using the essential SHAP features from each LOSO fold (i.e. 24 folds from 24 participants) across all cases. See Supplementary Note 3.4 for details on hyperparameter tuning.

Performance metrics comprising accuracy, F1-score, and area under the receiver operating characteristic (ROC) curve (AUC) were computed to assess the performance of the classifiers. This rigorous approach allowed us to explore the ability of the ML models to extract discriminative features from the HFA data, enabling optimal classification in the various cases.

### PCA Analysis

PCA was applied to the SHAP-identified contacts that overlapped across the three cases in each analysis. SL and RL used the trial-averaged HFA time courses in their respective windows, whereas ESR and CSR used their own fused data acquired after performing fusion, respectively. PCA was fitted to the combined data of the two classes, which were then projected onto the first three PCs. For visualisation purpose, trajectories were shifted to the origin at their initial point (SL, ESR, CSR) or final point (RL), which does not alter the metrics. Subclass comparisons followed the same steps. As PCA is linear, each trajectory splits exactly into per-contact contributions, giving paired values per contact for two-sided signflip permutation tests, with Holm correction across feature tests. A linear SVM was fitted in PCA space for visualisation, and its five-fold cross-validated accuracy is reported.

## Data Availability Statement

The analytical workflow was implemented using MATLAB 2024b on an Apple MacBook Pro with an M1 processor for preprocessing and visualisation, and Python 3.8 with Google Colab for machine learning and deep learning model development and deployment. Source code will be made available upon publication. The original dataset is openly available via the original study [20] (https://osf.io/9bzx8/).

## Acknowledgment

We thank Saugat Bhattacharyya, Brendan Lenfesty and Amin Azimi for helpful discussions. This work was supported by HSC R&D (STL/5540/19), MRC (MC OC 20020), and The Royal Society (IEC NSFC 252805).

## Competing interests

The authors declare no competing interests.

## Author contributions

U.K.N.M. and K.W-L. formulated the idea. U.K.N.M. and A.A. designed the experiments. U.K.N.M. performed and analysed the experiments. U.K.N.M. and K.W-L. analysed the data. S.G., R.G.OC., S.P.K and S.B. collated the data. U.K.N.M. and K.W-L. wrote the original draft of the paper. All authors provided feedback on data analysis and reviewed the manuscript.

## Additional information

**Supplementary information** The online version contains supplementary material. **Correspondence and requests for materials** should be addressed to KongFatt Wong-Lin or Umesh Kumar Naik Mudavath.

## Notes

### Competing Interest Statement

The authors have declared no competing interest.

