## Supplementary Information for "Neural latent representation of implicit perceptual decision confidence signatures from brain-wide intracranial EEG"

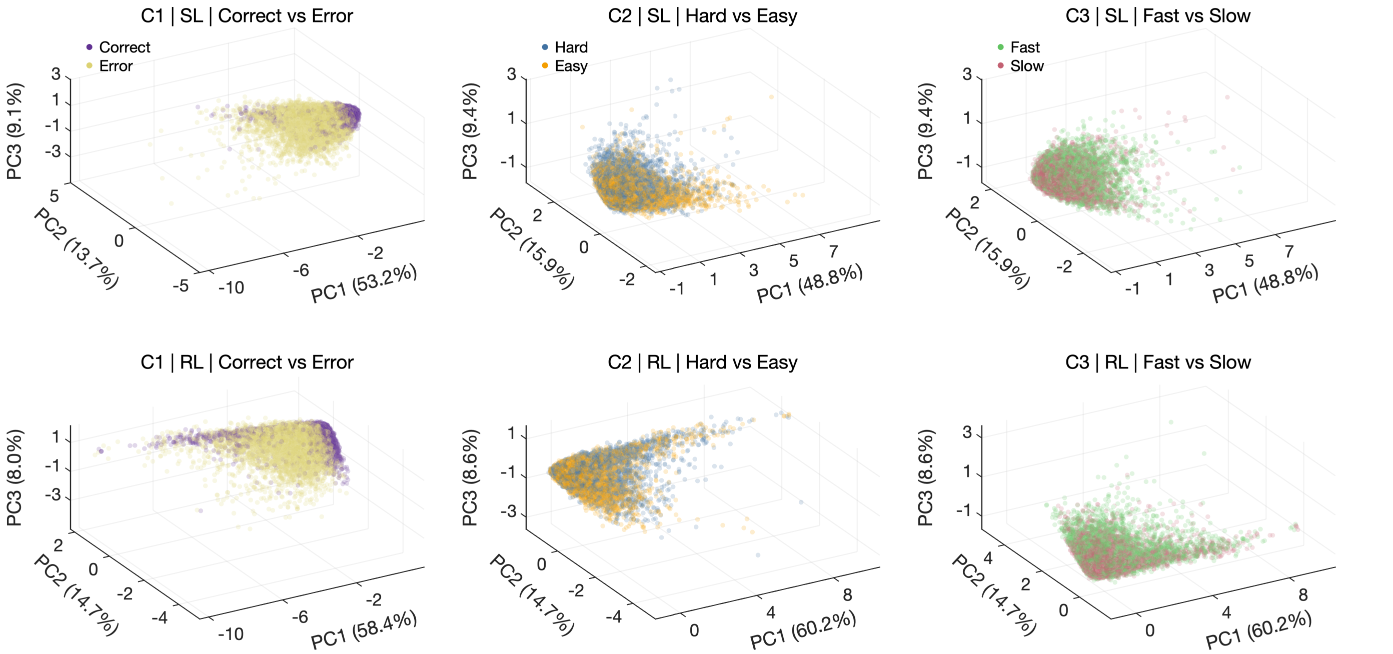


**Extended Data Fig. 1. PCA projections of the neural data in the manual response task.** Left, middle and right panels: Cases 1, 2 and 3, respectively. Top: PCA projections for the stimulus-locked (SL) analysis. Bottom: for the response-locked (RL) analysis. Each point represents the complete time course of one labelled neural (iEEG contact) observation, projected onto the top three principal components computed jointly across both classes within each panel. Points are not temporally aligned; their relative position reflects overall waveform similarity. Colours indicate the corresponding classes in each case.

**Supplementary Table 1. Reconstruction performance of adaptive transcoder for iEEG high-frequency activity signals during manual response task.**

| Case (Sub-condition) | Stimulus-Locked (SL) | | | | | Response-Locked (RL) | | | | |
|---|---|---|---|---|---|---|---|---|---|---|
| | MSE | RMSE | RRMSE | MAE | CC | MSE | RMSE | RRMSE | MAE | CC |
| 1 (Correct vs. error choice response) | 0.00018 | 0.01356 | 0.01525 | 0.01051 | 0.99839 | 0.00053 | 0.02301 | 0.02615 | 0.01527 | 0.99917 |
| 2 (Hard vs. easy task) | 0.00007 | 0.00818 | 0.00900 | 0.00487 | 0.99704 | 0.00015 | 0.01225 | 0.01363 | 0.00853 | 0.99838 |
| 3 (Fast vs. slow reaction time) | 0.00011 | 0.01061 | 0.01168 | 0.00837 | 0.99759 | 0.00007 | 0.00866 | 0.00963 | 0.00649 | 0.99852 |

MSE, mean squared error; RMSE, root mean squared error; RRMSE, relative root mean squared error; MAE, mean absolute error; CC, correlation coefficient.

**Supplementary Table 2. Performance of ensemble machine learning models with stratified 10-fold cross-validation for stimulus-locked manual response task.**

| Models | Case 1 | | | Case 2 | | | Case 3 | | |
|---|---|---|---|---|---|---|---|---|---|
| | Accuracy | F1-score | AUC | Accuracy | F1-score | AUC | Accuracy | F1-score | AUC |
| RFC | $0.88 \pm 0.00$ | $0.88 \pm 0.00$ | $0.94 \pm 0.00$ | $0.55 \pm 0.00$ | $0.55 \pm 0.00$ | $0.58 \pm 0.01$ | $0.57 \pm 0.00$ | $0.58 \pm 0.00$ | $0.61 \pm 0.00$ |
| AdaBoost | $0.81 \pm 0.00$ | $0.80 \pm 0.00$ | $0.89 \pm 0.00$ | $0.53 \pm 0.01$ | $0.52 \pm 0.07$ | $0.55 \pm 0.01$ | $0.58 \pm 0.00$ | $0.61 \pm 0.00$ | $0.60 \pm 0.00$ |
| LGBM | $0.89 \pm 0.00$ | $0.89 \pm 0.00$ | $0.95 \pm 0.00$ | $0.55 \pm 0.00$ | $0.57 \pm 0.00$ | $0.57 \pm 0.00$ | $0.57 \pm 0.00$ | $0.59 \pm 0.00$ | $0.60 \pm 0.01$ |
| XGB | $0.89 \pm 0.00$ | $0.88 \pm 0.00$ | $0.95 \pm 0.00$ | $0.53 \pm 0.01$ | $0.54 \pm 0.01$ | $0.56 \pm 0.01$ | $0.56 \pm 0.00$ | $0.57 \pm 0.00$ | $0.58 \pm 0.00$ |
| SVEC | $0.89 \pm 0.00$ | $0.88 \pm 0.00$ | $0.95 \pm 0.00$ | $0.55 \pm 0.00$ | $0.57 \pm 0.01$ | $0.58 \pm 0.01$ | $0.58 \pm 0.00$ | $0.60 \pm 0.01$ | $0.61 \pm 0.00$ |

SL, stimulus-locked; RFC, random forest classifier; AdaBoost, adaptive boosting; LGBM, light gradient boosting machine; XGB, extreme gradient boosting; SVEC, soft voting ensemble classifier; AUC, area under the ROC curve.

**Supplementary Table 3. Overall performance of different training strategies employing the LGBM classifier for manual response task.**

| Analysis | Case | CV | Without ATC | | | With ATC | | | Gherman et al. (2024) | | |
|---|---|---|---|---|---|---|---|---|---|---|---|
| | | | Accuracy | F1-score | AUC | Accuracy | F1-score | AUC | Accuracy | F1-score | AUC |
| | | LOSO | 0.89 ± 0.07 | 0.89 ± 0.08 | 0.96 ± 0.04 | 0.91 ± 0.08 | 0.90 ± 0.08 | 0.97 ± 0.03 | 0.82 ± 0.12 | 0.82 ± 0.12 | 0.90 ± 0.08 |
| | 1 | SGKF | 0.88 ± 0.06 | 0.88 ± 0.07 | 0.95 ± 0.03 | 0.90 ± 0.06 | 0.90 ± 0.07 | 0.97 ± 0.03 | 0.77 ± 0.06 | 0.77 ± 0.07 | 0.87 ± 0.06 |
| | | SKF | 0.89 ± 0.01 | 0.89 ± 0.01 | 0.95 ± 0.00 | 0.90 ± 0.01 | 0.90 ± 0.01 | 0.96 ± 0.01 | 0.81 ± 0.03 | 0.81 ± 0.03 | 0.87 ± 0.03 |
| | | LOSO | 0.53 ± 0.03 | 0.53 ± 0.04 | 0.55 ± 0.05 | 0.54 ± 0.04 | 0.53 ± 0.04 | 0.56 ± 0.05 | 0.56 ± 0.09 | 0.54 ± 0.12 | 0.63 ± 0.16 |
| SL | 2 | SGKF | 0.53 ± 0.01 | 0.53 ± 0.01 | 0.55 ± 0.02 | 0.54 ± 0.02 | 0.53 ± 0.02 | 0.56 ± 0.02 | 0.58 ± 0.06 | 0.57 ± 0.07 | 0.63 ± 0.08 |
| | | SKF | 0.55 ± 0.01 | 0.55 ± 0.01 | 0.58 ± 0.01 | 0.56 ± 0.02 | 0.56 ± 0.02 | 0.59 ± 0.02 | 0.56 ± 0.06 | 0.56 ± 0.06 | 0.58 ± 0.05 |
| | | LOSO | 0.55 ± 0.09 | 0.54 ± 0.09 | 0.57 ± 0.12 | 0.55 ± 0.10 | 0.55 ± 0.10 | 0.57 ± 0.13 | 0.57 ± 0.11 | 0.55 ± 0.12 | 0.63 ± 0.15 |
| | 3 | SGKF | 0.55 ± 0.04 | 0.54 ± 0.05 | 0.56 ± 0.06 | 0.55 ± 0.05 | 0.54 ± 0.05 | 0.56 ± 0.07 | 0.55 ± 0.10 | 0.54 ± 0.11 | 0.58 ± 0.12 |
| | | SKF | 0.58 ± 0.01 | 0.58 ± 0.01 | 0.61 ± 0.01 | 0.58 ± 0.01 | 0.58 ± 0.01 | 0.61 ± 0.01 | 0.60 ± 0.05 | 0.60 ± 0.05 | 0.66 ± 0.04 |
| | | LOSO | 0.87 ± 0.07 | 0.87 ± 0.08 | 0.95 ± 0.04 | 0.90 ± 0.07 | 0.89 ± 0.08 | 0.97 ± 0.04 | 0.78 ± 0.12 | 0.78 ± 0.12 | 0.87 ± 0.12 |
| | 1 | SGKF | 0.87 ± 0.06 | 0.87 ± 0.06 | 0.95 ± 0.03 | 0.89 ± 0.06 | 0.89 ± 0.07 | 0.96 ± 0.03 | 0.78 ± 0.07 | 0.77 ± 0.08 | 0.86 ± 0.06 |
| | | SKF | 0.87 ± 0.01 | 0.87 ± 0.01 | 0.94 ± 0.01 | 0.89 ± 0.01 | 0.89 ± 0.01 | 0.96 ± 0.01 | 0.79 ± 0.06 | 0.79 ± 0.06 | 0.86 ± 0.04 |
| | | LOSO | 0.53 ± 0.02 | 0.53 ± 0.02 | 0.55 ± 0.03 | 0.53 ± 0.03 | 0.53 ± 0.03 | 0.55 ± 0.04 | 0.51 ± 0.14 | 0.49 ± 0.15 | 0.52 ± 0.16 |
| RL | 2 | SGKF | 0.53 ± 0.02 | 0.53 ± 0.02 | 0.54 ± 0.03 | 0.53 ± 0.02 | 0.53 ± 0.02 | 0.55 ± 0.03 | 0.53 ± 0.08 | 0.53 ± 0.08 | 0.54 ± 0.11 |
| | | SKF | 0.55 ± 0.02 | 0.55 ± 0.02 | 0.58 ± 0.01 | 0.56 ± 0.01 | 0.56 ± 0.01 | 0.59 ± 0.02 | 0.56 ± 0.05 | 0.56 ± 0.05 | 0.58 ± 0.04 |
| | | LOSO | 0.55 ± 0.09 | 0.54 ± 0.09 | 0.57 ± 0.12 | 0.55 ± 0.09 | 0.54 ± 0.09 | 0.57 ± 0.13 | 0.57 ± 0.10 | 0.56 ± 0.10 | 0.62 ± 0.12 |
| | 3 | SGKF | 0.54 ± 0.05 | 0.54 ± 0.05 | 0.56 ± 0.06 | 0.55 ± 0.05 | 0.54 ± 0.05 | 0.56 ± 0.07 | 0.54 ± 0.04 | 0.54 ± 0.04 | 0.59 ± 0.06 |
| | | SKF | 0.57 ± 0.01 | 0.56 ± 0.01 | 0.60 ± 0.01 | 0.58 ± 0.01 | 0.58 ± 0.01 | 0.61 ± 0.02 | 0.58 ± 0.04 | 0.57 ± 0.04 | 0.63 ± 0.05 |
| | | LOSO | 0.91 ± 0.08 | 0.91 ± 0.08 | 0.97 ± 0.03 | 0.92 ± 0.08 | 0.92 ± 0.09 | 0.98 ± 0.03 | 0.85 ± 0.11 | 0.85 ± 0.12 | 0.95 ± 0.05 |
| | 1 | SGKF | 0.90 ± 0.07 | 0.90 ± 0.07 | 0.97 ± 0.03 | 0.92 ± 0.07 | 0.92 ± 0.09 | 0.98 ± 0.02 | 0.79 ± 0.08 | 0.78 ± 0.10 | 0.91 ± 0.05 |
| | | SKF | 0.91 ± 0.01 | 0.91 ± 0.01 | 0.97 ± 0.00 | 0.92 ± 0.01 | 0.92 ± 0.01 | 0.97 ± 0.00 | 0.81 ± 0.03 | 0.81 ± 0.03 | 0.90 ± 0.03 |
| | | LOSO | 0.53 ± 0.03 | 0.53 ± 0.04 | 0.56 ± 0.05 | 0.54 ± 0.04 | 0.54 ± 0.04 | 0.57 ± 0.05 | 0.59 ± 0.15 | 0.57 ± 0.15 | 0.59 ± 0.19 |
| ESR | 2 | SGKF | 0.53 ± 0.02 | 0.53 ± 0.02 | 0.55 ± 0.03 | 0.54 ± 0.02 | 0.54 ± 0.02 | 0.57 ± 0.03 | 0.60 ± 0.06 | 0.59 ± 0.07 | 0.63 ± 0.09 |
| | | SKF | 0.57 ± 0.01 | 0.57 ± 0.01 | 0.61 ± 0.02 | 0.58 ± 0.01 | 0.58 ± 0.01 | 0.63 ± 0.01 | 0.60 ± 0.05 | 0.59 ± 0.05 | 0.66 ± 0.05 |
| | | LOSO | 0.56 ± 0.10 | 0.55 ± 0.10 | 0.58 ± 0.13 | 0.56 ± 0.10 | 0.55 ± 0.10 | 0.59 ± 0.14 | 0.63 ± 0.15 | 0.62 ± 0.15 | 0.70 ± 0.17 |
| | 3 | SGKF | 0.55 ± 0.04 | 0.55 ± 0.04 | 0.57 ± 0.06 | 0.56 ± 0.05 | 0.55 ± 0.05 | 0.59 ± 0.07 | 0.59 ± 0.09 | 0.58 ± 0.09 | 0.63 ± 0.11 |
| | | SKF | 0.60 ± 0.01 | 0.60 ± 0.01 | 0.64 ± 0.01 | 0.61 ± 0.01 | 0.61 ± 0.01 | 0.65 ± 0.01 | 0.64 ± 0.03 | 0.64 ± 0.03 | 0.69 ± 0.04 |
| | | LOSO | 0.88 ± 0.07 | 0.88 ± 0.07 | 0.95 ± 0.04 | 0.93 ± 0.08 | 0.92 ± 0.08 | 0.98 ± 0.02 | 0.92 ± 0.09 | 0.92 ± 0.09 | 0.98 ± 0.03 |
| | 1 | SGKF | 0.87 ± 0.06 | 0.87 ± 0.06 | 0.94 ± 0.03 | 0.92 ± 0.07 | 0.92 ± 0.07 | 0.98 ± 0.02 | 0.88 ± 0.08 | 0.88 ± 0.09 | 0.96 ± 0.04 |
| | | SKF | 0.88 ± 0.00 | 0.88 ± 0.00 | 0.94 ± 0.00 | 0.92 ± 0.00 | 0.92 ± 0.00 | 0.97 ± 0.00 | 0.87 ± 0.04 | 0.87 ± 0.04 | 0.94 ± 0.03 |
| | | LOSO | 0.52 ± 0.02 | 0.52 ± 0.02 | 0.53 ± 0.02 | 0.53 ± 0.04 | 0.53 ± 0.04 | 0.56 ± 0.05 | 0.58 ± 0.10 | 0.57 ± 0.10 | 0.62 ± 0.17 |
| CSR | 2 | SGKF | 0.51 ± 0.01 | 0.51 ± 0.01 | 0.53 ± 0.01 | 0.54 ± 0.01 | 0.53 ± 0.01 | 0.56 ± 0.02 | 0.60 ± 0.05 | 0.59 ± 0.06 | 0.65 ± 0.05 |
| | | SKF | 0.53 ± 0.01 | 0.53 ± 0.01 | 0.55 ± 0.00 | 0.56 ± 0.00 | 0.56 ± 0.00 | 0.61 ± 0.01 | 0.63 ± 0.04 | 0.63 ± 0.04 | 0.68 ± 0.05 |
| | | LOSO | 0.53 ± 0.05 | 0.53 ± 0.05 | 0.55 ± 0.06 | 0.55 ± 0.12 | 0.55 ± 0.12 | 0.58 ± 0.16 | 0.59 ± 0.13 | 0.58 ± 0.13 | 0.61 ± 0.15 |
| | 3 | SGKF | 0.53 ± 0.03 | 0.53 ± 0.03 | 0.55 ± 0.04 | 0.54 ± 0.06 | 0.54 ± 0.06 | 0.57 ± 0.08 | 0.57 ± 0.08 | 0.57 ± 0.08 | 0.60 ± 0.10 |
| | | SKF | 0.55 ± 0.01 | 0.55 ± 0.01 | 0.57 ± 0.01 | 0.59 ± 0.01 | 0.59 ± 0.01 | 0.63 ± 0.01 | 0.62 ± 0.06 | 0.62 ± 0.07 | 0.67 ± 0.05 |

ATC, adaptive transcoder; SL, stimulus-locked; RL, response-locked; ESR, early SL-RL fusion; CSR, contrastive SL-RL fusion; CV, cross validation; LOSO, leave-one-subject-out; SGKF, stratified group $k$-fold; SKF, stratified $k$-fold; AUC, area under the ROC curve.

**Supplementary Table 4. Effect of SHAP-based feature-selection fraction on LGBM classifier with stratified group 5-fold cross-validation for manual response task.**

| Analysis | Case | 2% SHAP | | | 5% SHAP | | | 10% SHAP | | |
|---|---|---|---|---|---|---|---|---|---|---|
| | | Accuracy | F1-score | AUC | Accuracy | F1-score | AUC | Accuracy | F1-score | AUC |
| | 1 | 0.86 ± 0.03 | 0.86 ± 0.04 | 0.95 ± 0.02 | 0.89 ± 0.05 | 0.88 ± 0.06 | 0.95 ± 0.03 | 0.90 ± 0.04 | 0.89 ± 0.05 | 0.96 ± 0.02 |
| SL | 2 | 0.60 ± 0.03 | 0.60 ± 0.04 | 0.64 ± 0.04 | 0.58 ± 0.02 | 0.58 ± 0.03 | 0.63 ± 0.02 | 0.58 ± 0.01 | 0.59 ± 0.02 | 0.63 ± 0.02 |
| | 3 | 0.58 ± 0.04 | 0.58 ± 0.03 | 0.61 ± 0.04 | 0.59 ± 0.02 | 0.60 ± 0.03 | 0.64 ± 0.03 | 0.60 ± 0.02 | 0.60 ± 0.04 | 0.63 ± 0.02 |
| | 1 | 0.88 ± 0.03 | 0.88 ± 0.03 | 0.94 ± 0.03 | 0.88 ± 0.04 | 0.88 ± 0.05 | 0.94 ± 0.02 | 0.88 ± 0.05 | 0.87 ± 0.06 | 0.95 ± 0.03 |
| RL | 2 | 0.56 ± 0.01 | 0.56 ± 0.04 | 0.60 ± 0.02 | 0.56 ± 0.02 | 0.56 ± 0.03 | 0.61 ± 0.02 | 0.56 ± 0.02 | 0.57 ± 0.01 | 0.60 ± 0.02 |
| | 3 | 0.62 ± 0.03 | 0.62 ± 0.03 | 0.67 ± 0.03 | 0.62 ± 0.02 | 0.63 ± 0.03 | 0.67 ± 0.02 | 0.62 ± 0.02 | 0.64 ± 0.02 | 0.66 ± 0.04 |
| | 1 | 0.88 ± 0.04 | 0.88 ± 0.04 | 0.95 ± 0.02 | 0.84 ± 0.05 | 0.84 ± 0.05 | 0.92 ± 0.03 | 0.91 ± 0.04 | 0.91 ± 0.04 | 0.96 ± 0.02 |
| ESR | 2 | 0.59 ± 0.04 | 0.58 ± 0.04 | 0.63 ± 0.03 | 0.59 ± 0.01 | 0.58 ± 0.03 | 0.63 ± 0.02 | 0.58 ± 0.01 | 0.58 ± 0.02 | 0.62 ± 0.02 |
| | 3 | 0.64 ± 0.04 | 0.65 ± 0.06 | 0.69 ± 0.04 | 0.62 ± 0.04 | 0.62 ± 0.08 | 0.66 ± 0.04 | 0.62 ± 0.04 | 0.64 ± 0.04 | 0.67 ± 0.05 |
| | 1 | 0.89 ± 0.05 | 0.88 ± 0.07 | 0.95 ± 0.03 | 0.91 ± 0.06 | 0.91 ± 0.07 | 0.96 ± 0.04 | 0.91 ± 0.07 | 0.91 ± 0.08 | 0.96 ± 0.03 |
| CSR | 2 | 0.57 ± 0.03 | 0.58 ± 0.05 | 0.63 ± 0.03 | 0.57 ± 0.03 | 0.57 ± 0.04 | 0.62 ± 0.03 | 0.58 ± 0.02 | 0.59 ± 0.02 | 0.62 ± 0.03 |
| | 3 | 0.65 ± 0.05 | 0.66 ± 0.06 | 0.70 ± 0.05 | 0.66 ± 0.05 | 0.66 ± 0.09 | 0.71 ± 0.04 | 0.63 ± 0.04 | 0.63 ± 0.07 | 0.68 ± 0.04 |

SL, stimulus-locked; RL, response-locked; ESR, early SL-RL fusion; CSR, contrastive SL-RL fusion; AUC, area under the ROC curve.

**Supplementary Table 5. Comparison of all-contact, SHAP-identified, and Gherman et al. (2024) feature performance employing the LGBM classifier with stratified group 5-fold cross-validation for manual response task.**

| Analysis | Case | All iEEG HFA | | | 2% SHAP | | | Gherman et al. (2024) | | |
|---|---|---|---|---|---|---|---|---|---|---|
| | | Accuracy | F1-score | AUC | Accuracy | F1-score | AUC | Accuracy | F1-score | AUC |
| SL | 1 | 0.88 ± 0.06 | 0.88 ± 0.06 | 0.95 ± 0.03 | 0.86 ± 0.03 | 0.86 ± 0.04 | 0.95 ± 0.02 | 0.77 ± 0.06 | 0.77 ± 0.07 | 0.87 ± 0.06 |
| | 2 | 0.52 ± 0.01 | 0.52 ± 0.01 | 0.54 ± 0.01 | 0.60 ± 0.03 | 0.60 ± 0.04 | 0.64 ± 0.04 | 0.58 ± 0.06 | 0.57 ± 0.07 | 0.63 ± 0.08 |
| | 3 | 0.54 ± 0.04 | 0.54 ± 0.04 | 0.56 ± 0.06 | 0.58 ± 0.04 | 0.58 ± 0.03 | 0.61 ± 0.04 | 0.55 ± 0.10 | 0.54 ± 0.11 | 0.58 ± 0.12 |
| RL | 1 | 0.87 ± 0.05 | 0.86 ± 0.06 | 0.94 ± 0.03 | 0.88 ± 0.03 | 0.88 ± 0.03 | 0.94 ± 0.03 | 0.78 ± 0.07 | 0.77 ± 0.08 | 0.86 ± 0.06 |
| | 2 | 0.52 ± 0.01 | 0.52 ± 0.01 | 0.54 ± 0.02 | 0.56 ± 0.01 | 0.56 ± 0.04 | 0.60 ± 0.02 | 0.53 ± 0.08 | 0.53 ± 0.08 | 0.54 ± 0.11 |
| | 3 | 0.53 ± 0.04 | 0.53 ± 0.04 | 0.55 ± 0.06 | 0.62 ± 0.03 | 0.62 ± 0.03 | 0.67 ± 0.03 | 0.54 ± 0.04 | 0.54 ± 0.04 | 0.59 ± 0.06 |
| ESR | 1 | 0.91 ± 0.07 | 0.91 ± 0.07 | 0.97 ± 0.02 | 0.88 ± 0.04 | 0.88 ± 0.04 | 0.95 ± 0.02 | 0.79 ± 0.08 | 0.78 ± 0.10 | 0.91 ± 0.05 |
| | 2 | 0.54 ± 0.02 | 0.54 ± 0.02 | 0.56 ± 0.02 | 0.59 ± 0.04 | 0.58 ± 0.04 | 0.63 ± 0.03 | 0.60 ± 0.06 | 0.59 ± 0.07 | 0.63 ± 0.09 |
| | 3 | 0.55 ± 0.04 | 0.55 ± 0.04 | 0.58 ± 0.07 | 0.64 ± 0.04 | 0.65 ± 0.06 | 0.69 ± 0.04 | 0.59 ± 0.09 | 0.58 ± 0.09 | 0.63 ± 0.11 |
| CSR | 1 | 0.92 ± 0.07 | 0.92 ± 0.07 | 0.98 ± 0.02 | 0.89 ± 0.05 | 0.88 ± 0.07 | 0.95 ± 0.03 | 0.88 ± 0.08 | 0.88 ± 0.09 | 0.96 ± 0.04 |
| | 2 | 0.54 ± 0.01 | 0.53 ± 0.01 | 0.56 ± 0.02 | 0.57 ± 0.03 | 0.58 ± 0.05 | 0.63 ± 0.03 | 0.60 ± 0.05 | 0.59 ± 0.06 | 0.65 ± 0.05 |
| | 3 | 0.54 ± 0.06 | 0.54 ± 0.06 | 0.57 ± 0.08 | 0.65 ± 0.05 | 0.66 ± 0.06 | 0.70 ± 0.05 | 0.57 ± 0.08 | 0.57 ± 0.08 | 0.60 ± 0.10 |

SL, stimulus-locked; RL, response-locked; ESR, early SL-RL fusion; CSR, contrastive SL-RL fusion; AUC, area under the ROC curve.

**Supplementary Table 6. Classification performance employing the LGBM classifier with stratified group 5-fold cross-validation for vocal response task.**

| Analysis | Case | All iEEG HFA | | | 2% SHAP | | | Gherman et al. (2024) | | |
|---|---|---|---|---|---|---|---|---|---|---|
| | | Accuracy | F1-score | AUC | Accuracy | F1-score | AUC | Accuracy | F1-score | AUC |
| SL | 1 | 0.89 ± 0.06 | 0.89 ± 0.06 | 0.96 ± 0.03 | 0.90 ± 0.04 | 0.89 ± 0.03 | 0.98 ± 0.01 | 0.90 ± 0.07 | 0.88 ± 0.07 | 0.95 ± 0.04 |
| | 2 | 0.54 ± 0.06 | 0.53 ± 0.06 | 0.57 ± 0.10 | 0.58 ± 0.03 | 0.60 ± 0.06 | 0.67 ± 0.04 | 0.59 ± 0.04 | 0.56 ± 0.09 | 0.61 ± 0.09 |
| | 3 | 0.76 ± 0.17 | 0.74 ± 0.19 | 0.78 ± 0.20 | 0.77 ± 0.17 | 0.74 ± 0.22 | 0.80 ± 0.18 | 0.69 ± 0.12 | 0.65 ± 0.16 | 0.74 ± 0.15 |

SL, stimulus-locked; AUC, area under the ROC curve.

#### Supplementary Note 1

##### 1.1 Regional spatial distribution of SHAP-identified iEEG contacts

The region-wise spatial distribution of SHAP-identified iEEG contacts in Case 1 were distributed as follows: frontal (43.44%), parietal (22.81%), and temporal (20.62%) regions in SL analysis; frontal (36.14%), temporal (30.22%), and parietal (23.99%) regions in RL analysis; frontal (42.28%), temporal (24.07%), and parietal (22.53%) regions in ESR analysis; and frontal (44.31%), parietal (28.62%), and temporal (16.62%) regions in CSR analysis. While the corresponding distributions for Cases 2 and 3 are as follows. For Case 2, the distributions were: frontal (45.71%), parietal (24.76%), and temporal (19.37%) regions in SL analysis; parietal (37.20%), frontal (32.62%), and temporal (20.73%) regions in RL analysis; frontal (42.14%), parietal (25.79%), and temporal (23.27%) regions in ESR analysis; and frontal (44.59%), parietal (23.25%), and temporal (22.29%) regions in CSR analysis (Fig. 3A). In Case 3, the distributions were: frontal (43.49%), parietal (24.13%), and temporal (20.00%) regions in SL analysis; frontal (37.96%), parietal (32.72%), and temporal (23.15%) regions in RL analysis; frontal (44.34%), parietal (24.84%), and temporal (21.07%) regions in ESR analysis; and frontal (40.75%), parietal (29.47%), and temporal (20.06%) regions in CSR analysis (Fig. 3A).

Across the three cases, the distributions were: frontal (70.59%), temporal (17.65%), and parietal (11.76%) in SL analysis; temporal (38.89%), parietal (27.78%), and frontal (22.22%) in RL analysis; frontal (44.44%), temporal (33.33%), and occipital (22.22%) in ESR analysis; and frontal (58.82%), parietal (20.59%), and temporal (17.65%) in CSR analysis.

##### 1.2 Subregional spatial distribution of SHAP-identified iEEG contacts

At the subregional level, the top-ranked subregions for Case 1 were the lateral orbitofrontal, insula, and superior temporal regions in SL; superior temporal, middle temporal, and supramarginal regions in RL; superior temporal, lateral orbitofrontal, and inferior temporal regions in ESR; and precentral, postcentral, and supramarginal regions in CSR analysis (Fig. 3A). For Case 2, the top-ranked subregions were the precentral, postcentral, and lateral orbitofrontal in SL; postcentral, precentral, and supramarginal in RL; precentral, postcentral, and superior parietal in ESR; and precentral, superior frontal, and supramarginal in CSR analysis. For Case 3, the top-ranked subregions were precentral, superior parietal, and supramarginal in SL; precentral, postcentral, and supramarginal in RL; precentral, postcentral, and supramarginal in ESR; and supramarginal, precentral, and postcentral in CSR analysis (Fig. 3A).

Across the three cases, the shared contacts were mainly in the lateral orbitofrontal, medial orbitofrontal and inferior temporal regions in SL; superior temporal, supramarginal and inferior temporal regions in RL; precentral, fusiform and inferior temporal regions in ESR; precentral, pars opercularis and postcentral regions in CSR analysis.

##### 1.3 Overlap of SHAP-identified iEEG contacts

The largest overlap with effector-independent labels was observed for Case 2 (SL) and Case 2 (ESR) (both 21.4%), while the largest overlap with the effector labels was observed for Case 3 (RL) (28.8%), followed by Case 2 (RL) (26.6%), Case 1 (CSR) (26.2%), and Case 3 (ESR) (24.5%) (Fig. 3C). The subcase intersections also showed very limited overlaps with the labels reported by the previous study. The overlaps with the effector-independent labels ranged up to 5.5%, and the overlaps with the effector-dependent labels ranged from up to 2.6%. The highest overlap was observed in CSR, with 5.5% for the effector-independent labels and 2.6% for the effector-dependent labels, whereas the other analyses showed only marginal overlaps.

**Supplementary Table 7. Anatomical distribution of essential contacts identified for stimulus-locked analysis (indicative of effector-independent iEEG contacts).**

| Participant ID | Contact Location | Hemisphere | Brain Atlas Region | Brain Lobe | Detailed Anatomical Label |
|---|---|---|---|---|---|
| S2 | HDG58 | R | Supramarginal | Parietal | Supramarginal gyrus |
| S3 | RFo9 | R | Lateral Orbitofrontal | Frontal | Anterior segment of the circular sulcus of the insula |
| S5 | RFi11 | R | Pars Triangularis | Frontal | Horizontal ramus of the anterior segment of the lateral sulcus (or fissure) |
| S7 | RFo11 | R | Pars Triangularis | Frontal | Lateral orbital sulcus |
| S11 | RTx1 | R | Medial Orbitofrontal | Frontal | Straight gyrus, Gyrus rectus |
| | RTx2 | R | Medial Orbitofrontal | Frontal | Straight gyrus, Gyrus rectus |
| | RTx3 | R | Lateral Orbitofrontal | Frontal | Subcallosal area, subcallosal gyrus |
| | RTx4 | R | Lateral Orbitofrontal | Frontal | Orbital gyri |
| S12 | LFi7 | L | Lateral Orbitofrontal | Frontal | Anterior segment of the circular sulcus of the insula |
| | RFo12 | R | Pars Orbitalis | Frontal | Lateral orbital sulcus |
| S13 | LFs7 | L | Precentral | Frontal | Superior part of the precentral sulcus |
| | LTi11 | L | Inferior Temporal | Temporal | NaN |
| S15 | LDa5 | L | Unknown | Unknown | NaN |
| S20 | LDp10 | L | Superior Temporal | Temporal | Superior temporal sulcus (parallel sulcus) |
| | LTi3 | L | Inferior Temporal | Temporal | Inferior temporal gyrus (T3) |
| S21 | ROF5 | R | Medial Orbitofrontal | Frontal | Subcallosal area, subcallosal gyrus |
| | ROF7 | R | Medial Orbitofrontal | Frontal | Subcallosal area, subcallosal gyrus |
| S24 | LFp14 | L | Postcentral | Parietal | Postcentral gyrus |

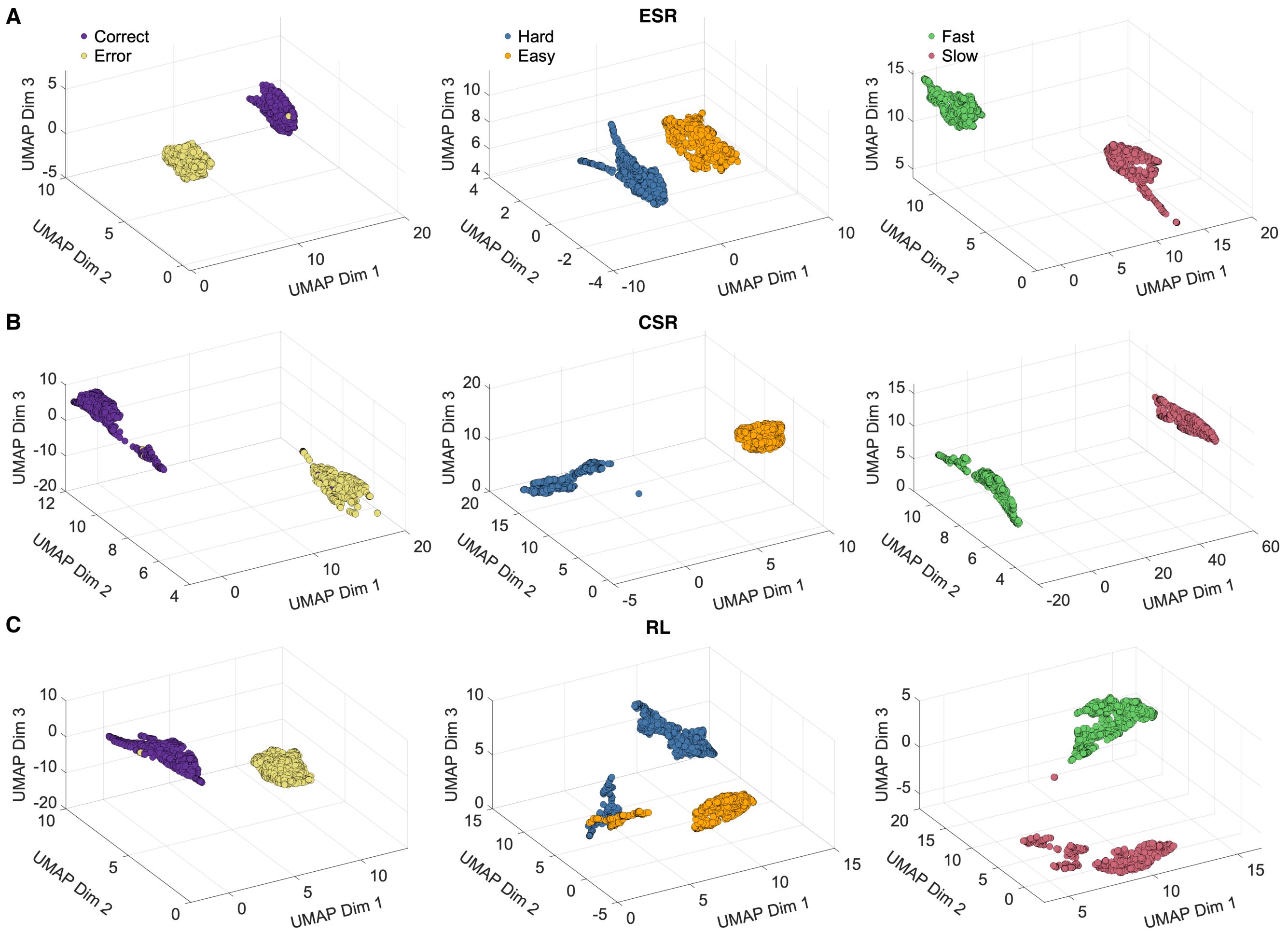


**Extended Data Fig. 2. Low-dimensional latent representations from adaptive transcoder under different analyses.** Left, middle and right panels: Cases 1, 2 and 3, respectively. **A-C.** 3D visualisations of UMAP projection in manual responses task for early SL-RL (ESR) fusion (**A**), contrastive SL-RL (CSR) fusion (**B**), and response-locked (RL) (**C**) analyses.

**Supplementary Table 8. Anatomical distribution of essential contacts identified for response-locked analysis (indicative of effector-independent iEEG contacts).**

| Participant ID | Contact Location | Hemisphere | Brain Atlas Region | Brain Lobe | Detailed Anatomical Label |
|---|---|---|---|---|---|
| S3 | RPl9 | R | Postcentral | Parietal | Postcentral sulcus |
| S4 | RDh1 | R | Fusiform | Temporal | Temporal pole |
| | RTi4 | R | Inferior Temporal | Temporal | Inferior temporal gyrus (T3) |
| S5 | RPc12 | R | Superior Parietal | Parietal | Postcentral sulcus |
| | RPi11 | R | Supramarginal | Parietal | Postcentral sulcus |
| S6 | RTx1 | R | Medial Orbitofrontal | Frontal | Subcallosal area, subcallosal gyrus |
| | RTx2 | R | Lateral Orbitofrontal | Frontal | Subcallosal area, subcallosal gyrus |
| S7 | RDa10 | R | Middle Temporal | Temporal | Middle temporal gyrus (T2) |
| S8 | RGrid220 | R | Supramarginal | Parietal | Supramarginal gyrus |
| S9 | LGrid40 | L | Supramarginal | Parietal | Supramarginal gyrus |
| S11 | RTx1 | R | Medial Orbitofrontal | Frontal | Straight gyrus, Gyrus rectus |
| S13 | LFp8 | L | Precentral | Frontal | Central sulcus (Rolando's fissure) |
| | LTx9 | L | Superior Temporal | Temporal | Planum polare of the superior temporal gyrus |
| S14 | ROl8 | R | Lateral Occipital | Occipital | Middle occipital gyrus (O2, lateral occipital gyrus) |
| S15 | LO3 | L | Cuneus | Occipital | Cuneus (O6) |
| S20 | LDh9 | L | Superior Temporal | Temporal | Superior temporal sulcus (parallel sulcus) |
| | LDp11 | L | Superior Temporal | Temporal | Lateral aspect of the superior temporal gyrus |
| S23 | RTo8 | R | Inferior Temporal | Temporal | Inferior temporal gyrus (T3) |

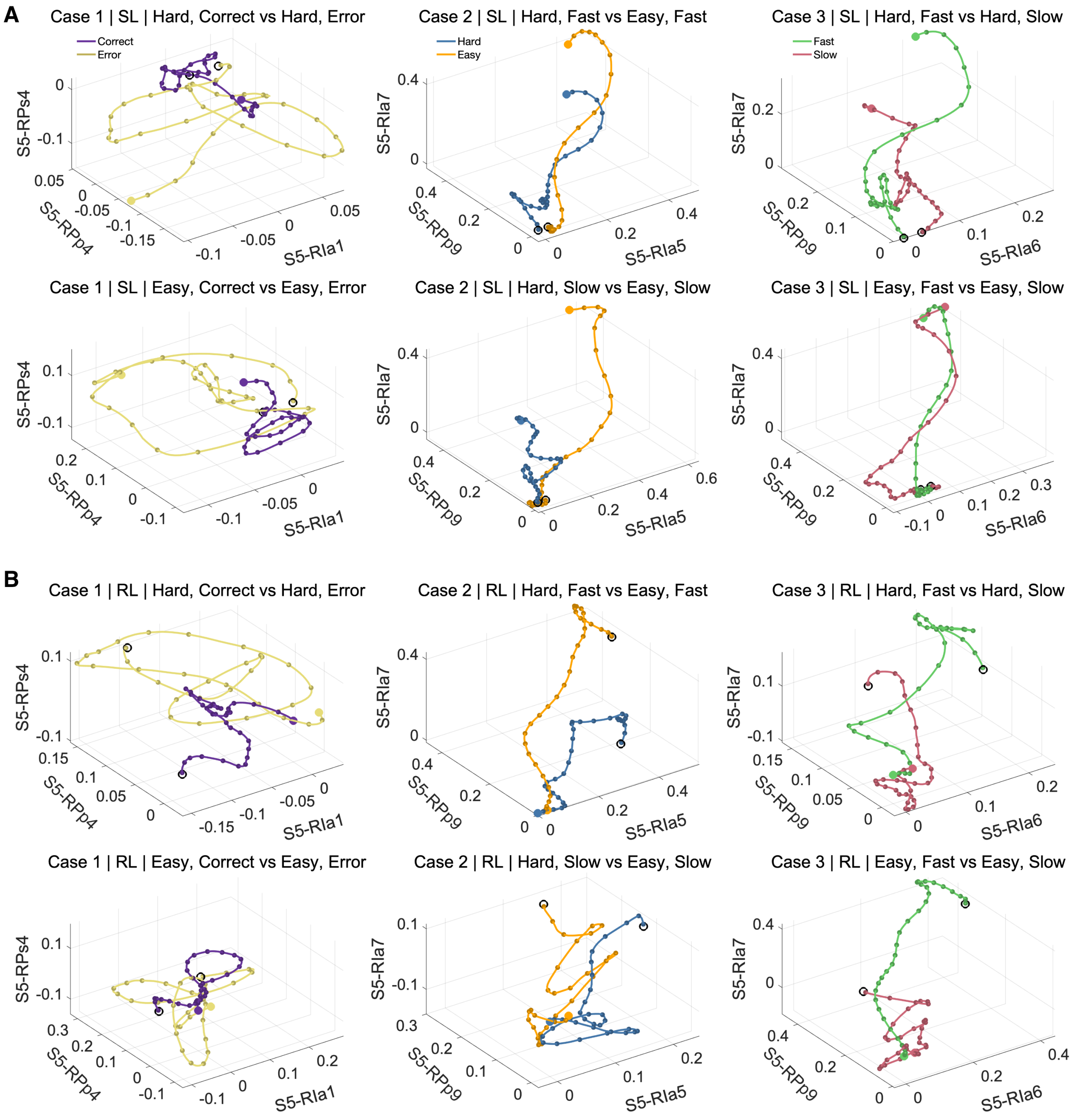


**Extended Data Fig. 3. Contact specific neural trajectories across different sub-conditions.** Left, middle and right panels: Cases 1, 2 and 3, respectively. **A.** Trial-averaged neural trajectories projected onto high-ranked feature dimensions for a sample participant for stimulus-locked (SL) analysis. Open circles: starting points; filled circles: end points. **B.** for response-locked (RL) analysis.

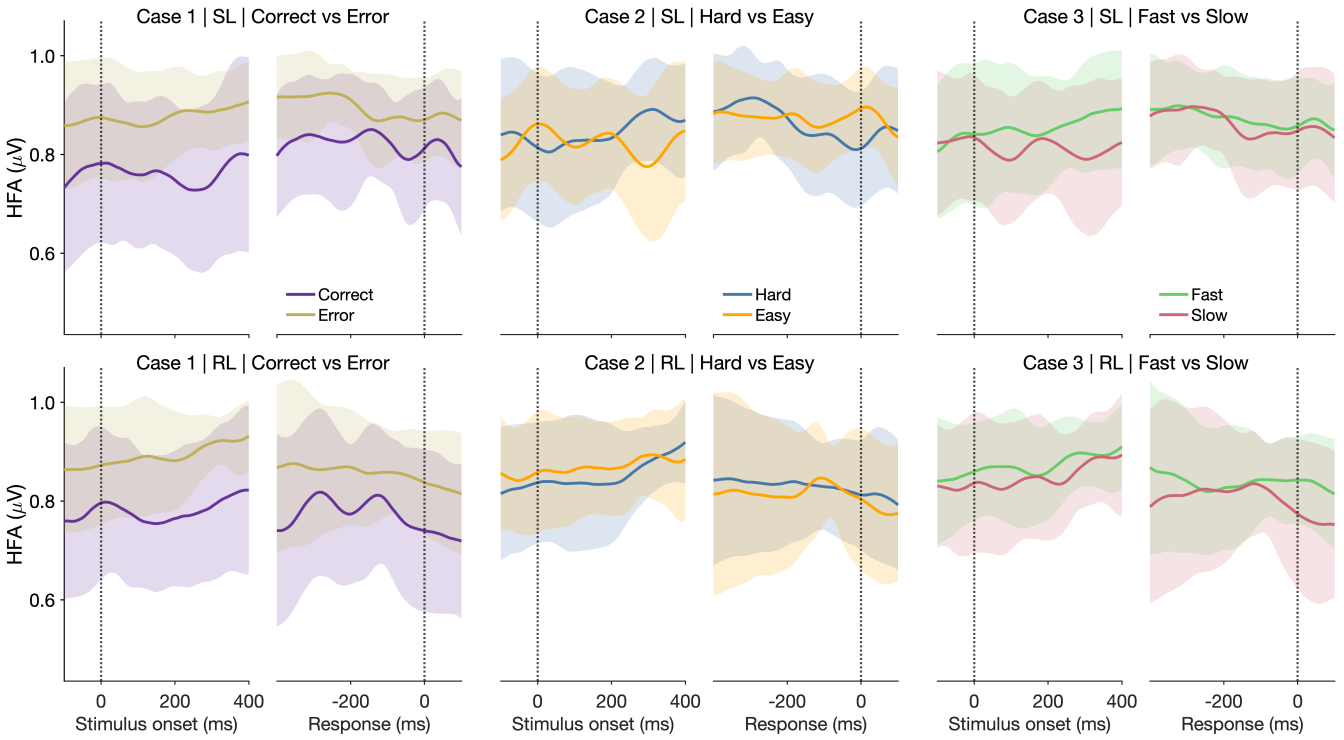


**Extended Data Fig. 4. Unweighted grand-averaged iEEG HFA time course for SHAP-identified contacts.** Left, middle and right panels: Cases 1, 2 and 3, respectively. Grand-averaged neural activity timecourse for SHAP-identified contacts under stimulus-locked (SL, top) and response-locked (RL, bottom) analyses without feature weighting.

**Supplementary Table 9. Anatomical distribution of essential contacts identified for early fusion analysis (indicative of effector-independent iEEG contacts).**

| Participant ID | Contact Location | Hemisphere | Brain Atlas Region | Brain Lobe | Detailed Anatomical Label |
|---|---|---|---|---|---|
| S3 | RPs7 | R | Precentral | Frontal | Central sulcus (Rolando's fissure) |
| | RPs8 | R | Precentral | Frontal | Precentral gyrus |
| S4 | RDh1 | R | Fusiform | Temporal | Temporal pole |
| | RTp9 | R | Inferior Temporal | Temporal | Inferior temporal sulcus |
| S11 | RDa6 | R | Unknown | Unknown | NaN |
| | RTx4 | R | Lateral Orbitofrontal | Frontal | Orbital gyri |
| S18 | LHi1 | L | Pericalcarine | Occipital | Occipital pole |
| S20 | LDp10 | L | Superior Temporal | Temporal | Superior temporal sulcus (parallel sulcus) |
| S21 | RMO12 | R | Lateral Occipital | Occipital | NaN |
| | ROF5 | R | Medial Orbitofrontal | Frontal | Subcallosal area, subcallosal gyrus |

**Supplementary Table 10. Anatomical distribution of essential contacts identified for contrastive fusion analysis (indicative of effector-independent iEEG contacts).**

| Participant ID | Contact Location | Hemisphere | Brain Atlas Region | Brain Lobe | Detailed Anatomical Label |
|---|---|---|---|---|---|
| S1 | RFi12 | R | Pars Opercularis | Frontal | Opercular part of the inferior frontal gyrus |
| S3 | LIp14 | L | Supramarginal | Parietal | NaN |
| | RFi7 | R | Rostral Middle Frontal | Frontal | Middle frontal gyrus (F2) |
| S4 | RDh5 | R | Unknown | Unknown | NaN |
| S5 | RFi11 | R | Pars Triangularis | Frontal | Horizontal ramus of the anterior segment of the lateral sulcus (or fissure) |
| | RIa4 | R | Pars Opercularis | Frontal | Opercular part of the inferior frontal gyrus |
| | RIa7 | R | Pars Opercularis | Frontal | Inferior part of the precentral sulcus |
| S6 | RTp7 | R | Fusiform | Temporal | Lateral occipito-temporal sulcus |
| | RTx2 | R | Lateral Orbitofrontal | Frontal | Subcallosal area, subcallosal gyrus |
| S7 | RDa10 | R | Middle Temporal | Temporal | Middle temporal gyrus (T2) |
| S9 | LFx4 | L | Rostral Middle Frontal | Frontal | NaN |
| S10 | LIp13 | L | Precentral | Frontal | Central sulcus (Rolando's fissure) |
| | LIp14 | L | Precentral | Frontal | Central sulcus (Rolando's fissure) |
| | LIp15 | L | Precentral | Frontal | Central sulcus (Rolando's fissure) |
| | RIp15 | R | Precentral | Frontal | Precentral gyrus |
| S11 | RTx1 | R | Medial Orbitofrontal | Frontal | Straight gyrus, Gyrus rectus |
| S12 | LFp8 | L | Precentral | Frontal | Central sulcus (Rolando's fissure) |
| S13 | LFp6 | L | Precentral | Frontal | Central sulcus (Rolando's fissure) |
| | LFs3 | L | Superior Frontal | Frontal | Superior frontal gyrus (F1) |
| | LFs7 | L | Precentral | Frontal | Superior part of the precentral sulcus |
| | RFo1 | R | Medial Orbitofrontal | Frontal | Straight gyrus, Gyrus rectus |
| S14 | RPa11 | R | Supramarginal | Parietal | Postcentral sulcus |
| | RPa7 | R | Postcentral | Parietal | Postcentral sulcus |
| S15 | LDa4 | L | Unknown | Unknown | NaN |
| | LIa6 | L | Pars Opercularis | Frontal | Opercular part of the inferior frontal gyrus |
| | LTo9 | L | Inferior Temporal | Temporal | Inferior temporal sulcus |
| S16 | LFp5 | L | Caudal Middle Frontal | Frontal | Superior frontal sulcus |
| | LPc8 | L | Supramarginal | Parietal | Postcentral sulcus |
| S18 | LDa15 | L | Lateral Occipital | Occipital | Anterior occipital sulcus and preoccipital notch (temporo-occipital incisure) |
| S19 | LFi11 | L | Pars Opercularis | Frontal | Opercular part of the inferior frontal gyrus |
| | LIp14 | L | Postcentral | Parietal | Postcentral sulcus |
| S20 | LDh9 | L | Superior Temporal | Temporal | Superior temporal sulcus (parallel sulcus) |
| | LTi1 | L | Inferior Temporal | Temporal | Inferior temporal gyrus (T3) |
| S23 | RDp8 | R | Banks of STS | Temporal | Superior temporal sulcus (parallel sulcus) |
| S24 | LFp12 | L | Postcentral | Parietal | Postcentral gyrus |
| | LPc5 | L | Superior Parietal | Parietal | NaN |

### Supplementary Note 2

#### *2.1 Robust low-dimensional neural latent representations for ESR and CSR fusion analyses*

SL and RL trajectory PCA results are reported in the main text (Fig. 5). Here, we extend the same analysis to the ESR and CSR fusion analyses. When we projected the grand-averaged neural trajectories for ESR and CSR analyses in PCA space, we again found that the first three principal components (PCs) sufficed to account for about 75% of the data variance (Extended Data Figs. 5C-D). Moreover, the neural trajectories were again well separable, with SVM classification accuracies of 93.8% (Case 1, ESR), 71.5% (Case 2, ESR), 73% (Case 3, ESR). 84.4% (Case 1, CSR), 61.7% (Case 2, CSR) and 54.4% (Case 3, CSR). We repeated the PCA analyses for various subcases and found qualitatively similar trajectory patterns within each analysis, though CSR required more PCs to capture comparable variance, both at the class level (Extended Data Fig. 5) and at the subclass level (Extended Data Fig. 10).

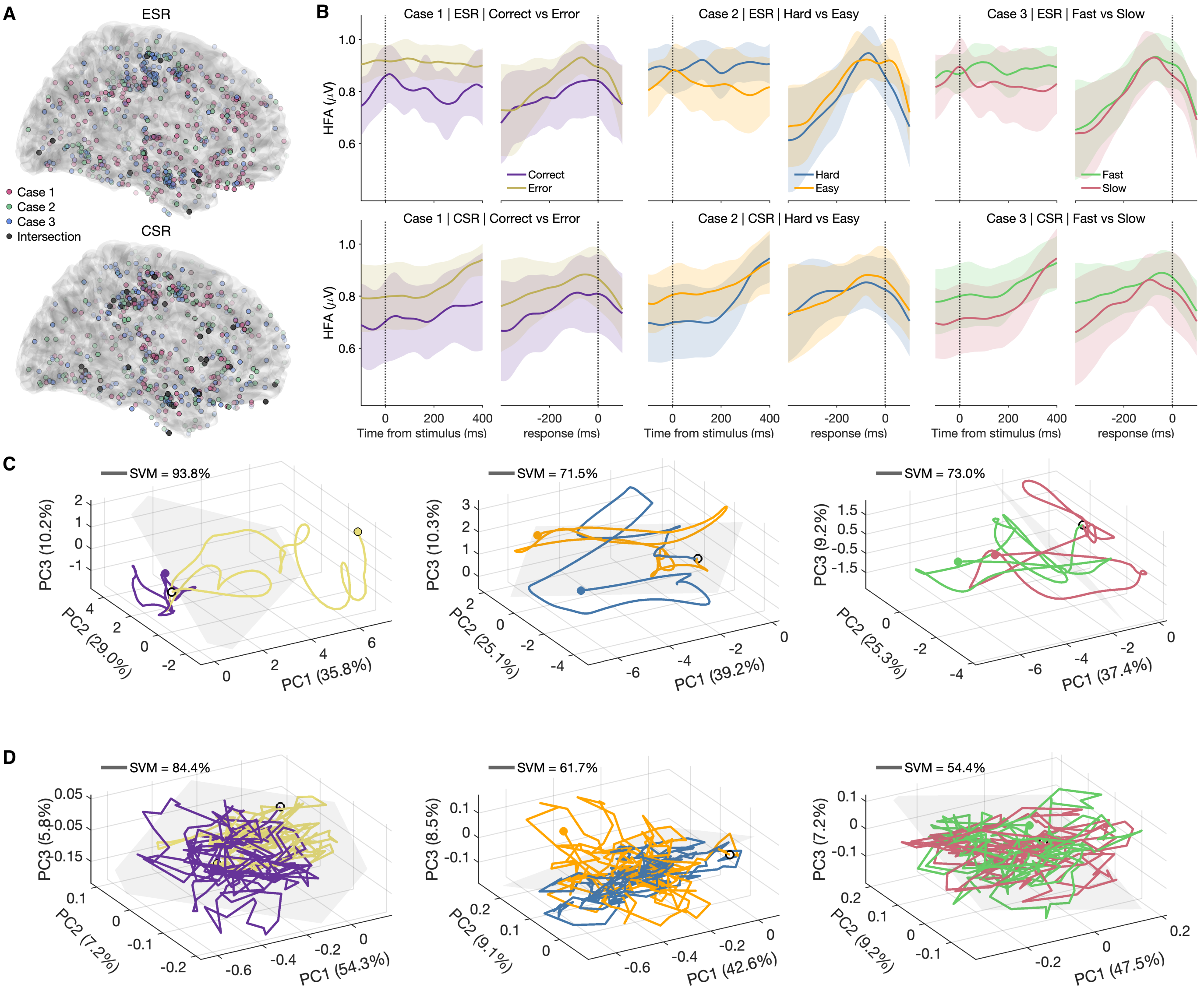


**Extended Data Fig. 5. SHAP-identified contact locations and grand-averaged activities for ESR and CSR analyses. A.** Dark grey: Projected structural MRI topographical plot mapping of essential HFA contacts (Case 1 (red) ∩ Case 2 (green) ∩ Case 3 (blue)) identified by SHAP in the ESR (top) and CSR (bottom) analyses. Grey: 10 (36) contacts overlapped across all cases for ESR (CSR). Colour transparency encodes SHAP importance (more opaque denotes higher importance). **B.** PCA-weighted grand-averaged neural activity timecourses; ESR (top) and CSR (bottom) analyses. **C–D.** Grand-averaged neural trajectories projected onto PCA space of the SHAP-identified top-ranked features that intersected (A, grey regions) across all three cases. **C,** ESR analysis; **D,** CSR analysis. Left to right: Cases 1, 2 and 3, respectively. Linear SVM decision hyperplanes (grey planes) overlaid for further classification of neural trajectories. Open circles: starting points; filled circles: end points. Colour labels as in B.

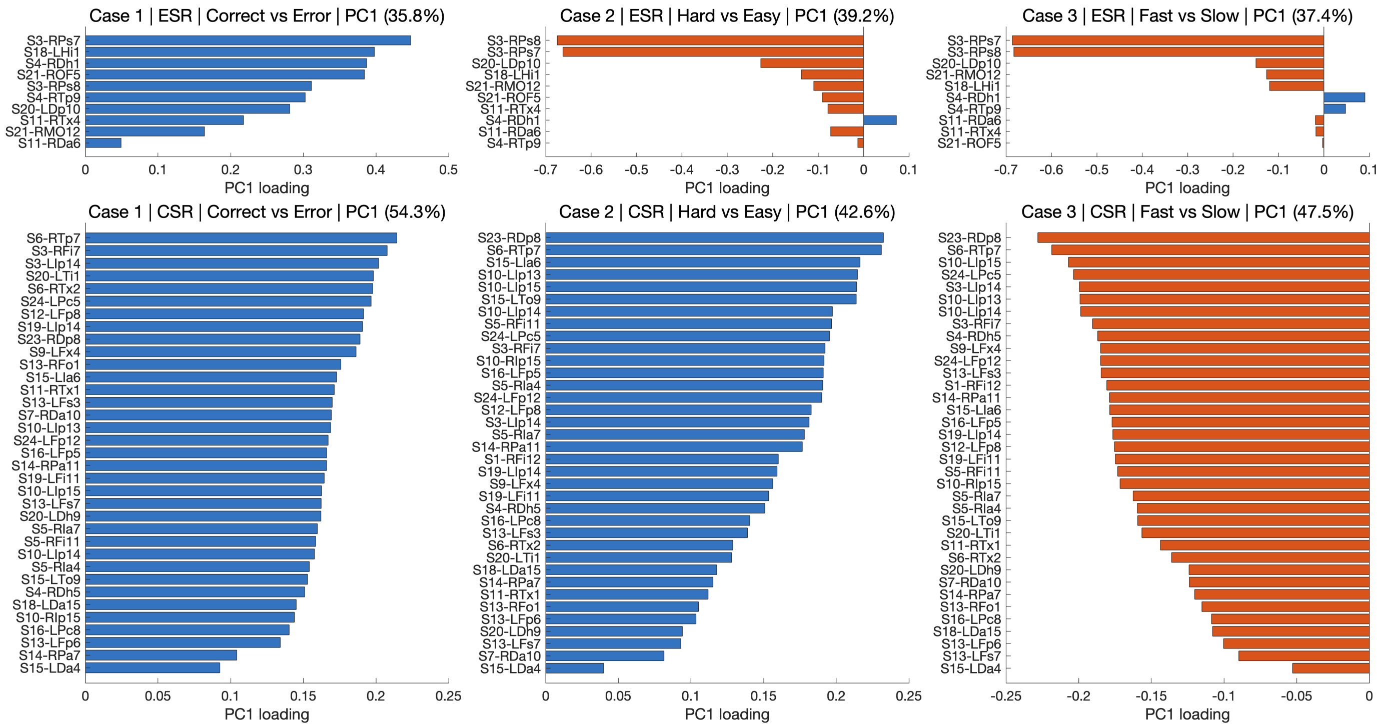


**Extended Data Fig. 6. PC1 loading distribution across fusion analyses.** PC1 loadings of essential iEEG contacts for early SL-RL (ESR, top) fusion and contrastive SL-RL (CSR, bottom) fusion analyses, computed from ESR and CSR data, respectively.

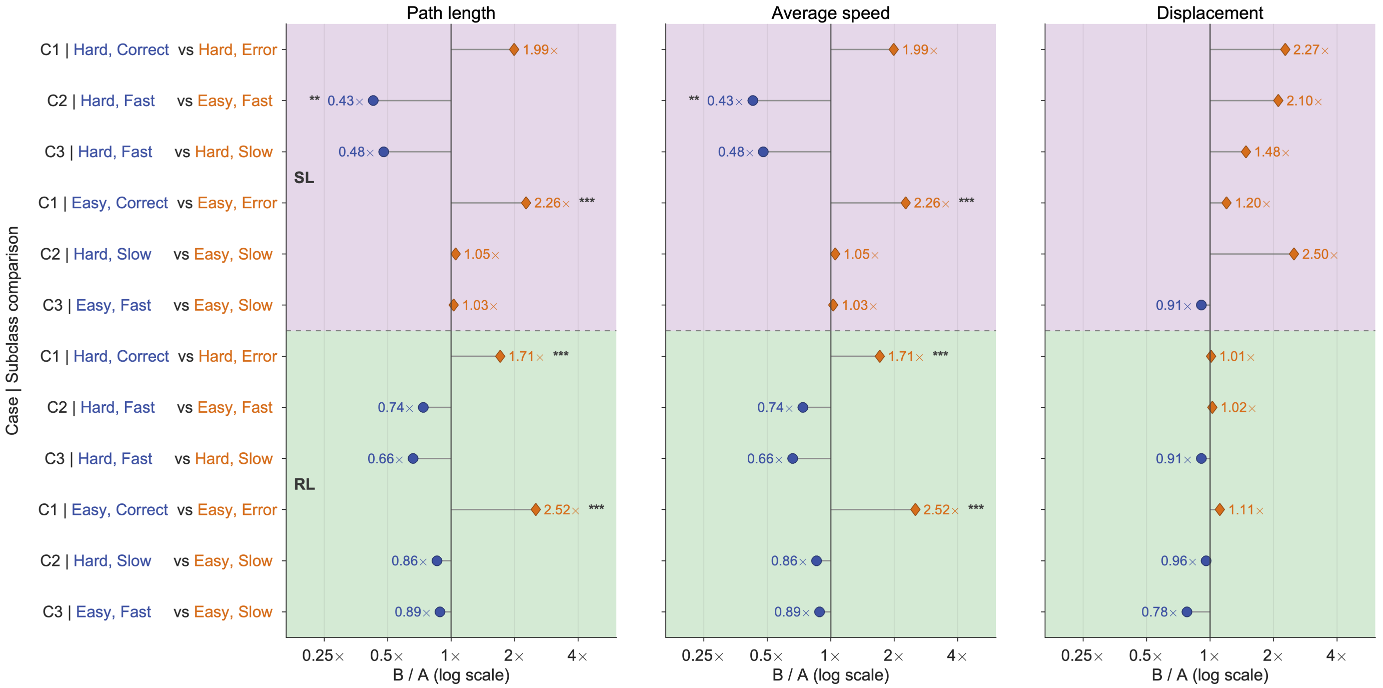


**Extended Data Fig. 7. Subclass-level PCA trajectory metrics for the SL and RL analyses.** Top: stimulus-locked (SL); bottom: response-locked (RL). Left to right: path length, average speed and displacement, respectively. The metrics were calculated from the grand-averaged PCA trajectories for each subclass comparison. Values shown beside the markers were calculated directly from the grand-averaged trajectories, whereas statistical significance was assessed across paired contacts using two-sided sign-flip permutation tests.

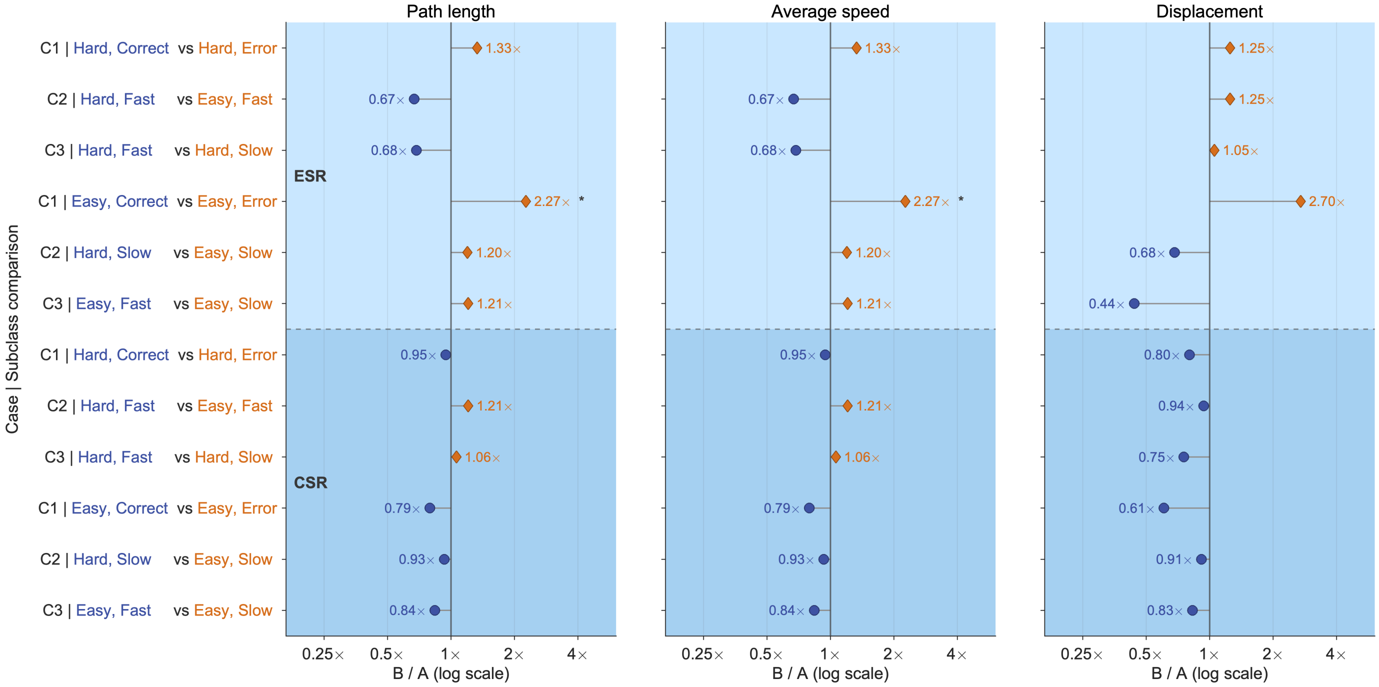


**Extended Data Fig. 8. Subclass-level PCA trajectory metrics for the ESR and CSR analyses.** Top: early SL-RL (ESR) fusion; bottom: contrastive SL-RL (CSR) fusion. Left to right: path length, average speed and displacement, respectively. The metrics were calculated from the grand-averaged PCA trajectories for each subclass comparison.

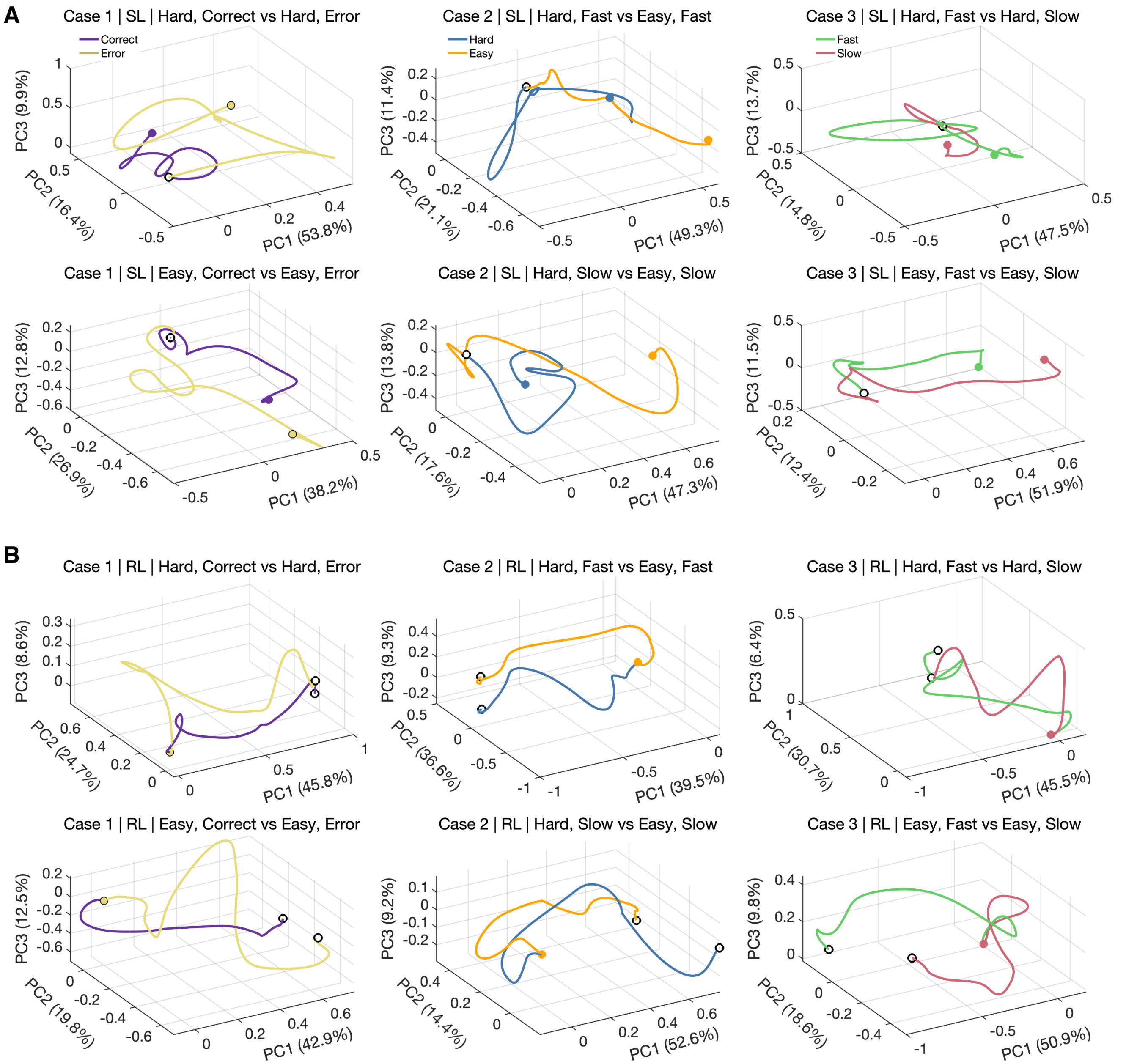


**Extended Data Fig. 9. Grand-averaged neural trajectories across different sub-conditions in PCA space.** Left, middle and right panels: Cases 1, 2 and 3, respectively. **A.** Stimulus-locked (SL) grand-averaged neural trajectories projected onto the first three principal components. Open circles: starting points; filled circles: end points. **B.** for response-locked (RL) analysis.

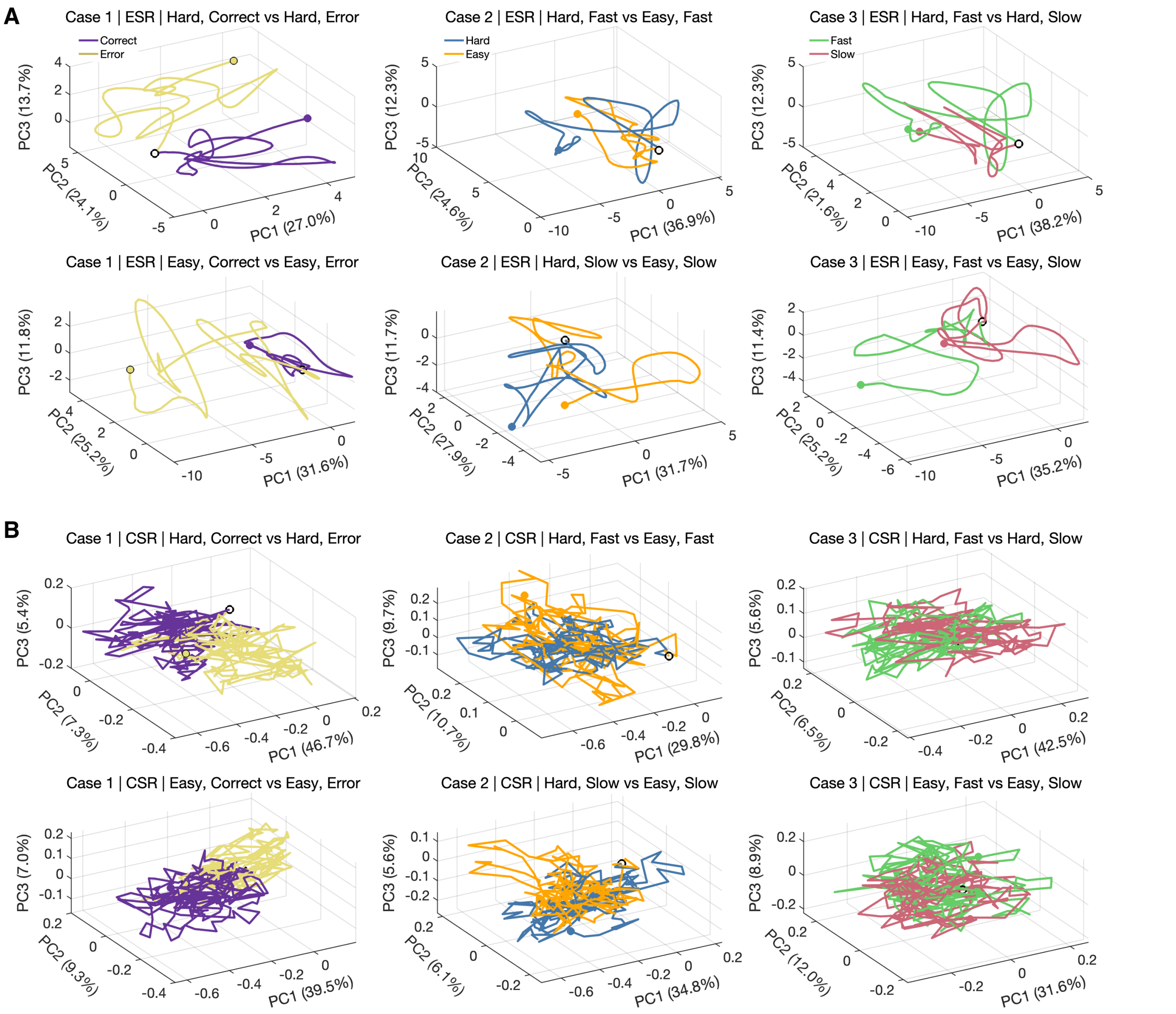


**Extended Data Fig. 10. Grand-averaged neural trajectories across different sub-conditions in PCA space.** Left, middle and right panels: Cases 1, 2 and 3, respectively. **A.** Early SL-RL (ESR) fusion grand-averaged neural trajectories projected onto the first three principal components. Open circles: starting points; filled circles: end points. **B.** for contrastive SL-RL (CSR) fusion analysis.

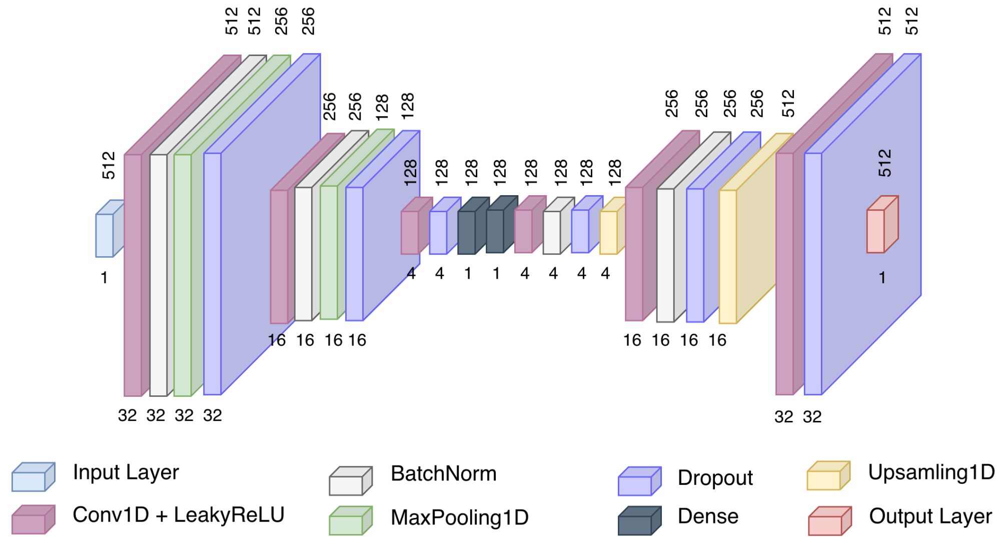


**Extended Data Fig. 11. The system architecture of the proposed autoencoder model framework.** Numbers in each layer: encoder comprises Conv1D layers with 32, 16, and 4 filters followed by a 1-unit bottleneck; decoder comprises a 1-unit dense layer, Conv1D layers with 4, 16, and 32 filters, and a final Conv1D layer with 1 filter. All convolutional layers use kernel size 32, dropout rate 0.1, and pooling/upsampling factor 2.

#### Supplementary Note 3

##### 3.1 Mathematical formulation of the proposed temporal convolutional autoencoder (TCA) framework

The proposed TCA framework can be mathematically formulated as follows. Let $i = \mathbf{I}^{S\times T\times 1}_{initial}$ represent the input HFA data, where $T$ corresponds to the number of time points in each HFA signal $\mathbf{I}$, and $S$ is the number of HFA samples. Moreover, E and D represent the encoder and decoder of the TCA. Then, the encoder produces a compressed (latent) representation given by, $j = \mathbf{Z}^{S\times T'\times 1}_{compressed}$, where $T'$ is the compressed temporal 1D representation. The decoder reconstructs the input, yielding $\hat{i} = \mathbf{I}^{S\times T\times 1}_{initial}$. These operations can simply be denoted as $j = E(i)$ and $\hat{i} = D(j)$. The reconstruction error $r^{TCA}$ can be used to evaluate the TCAs' performance, and is described by:

$$r^{TCA} = M_{TCA}\Big(\hat{i}^{(x)},\, i^{(x)}\Big) \tag{1}$$

where the $M_{TCA}$ function represents a measurement of difference, much like the ubiquitous square Euclidean distance, as follows:

$$M_{TCA}\Big(\hat{i}^{(x)},\, i^{(x)}\Big) = \frac{1}{2}\left\|\hat{i}^{(x)} - i^{(x)}\right\|^2 \tag{2}$$

while the cost function $N_{TCA}$ can be expressed in its generic form as:

$$N_{TCA} = \frac{1}{S}\sum_{x=1}^{S} L_{TCA}\Big(D\big(E(i^{(x)})\big),\, i^{(x)}\Big) \tag{3}$$

##### 3.2 Early SL-RL (ESR) fusion

For ESR, let $\mathbf{x}_{SL}$ and $\mathbf{x}_{RL}$ denote the SL and RL embeddings extracted independently for the same HFA sample. In ESR, we form a joint representation by feature-level concatenation, $\mathbf{x}_F = [\mathbf{x}_{SL}, \mathbf{x}_{RL}]$ (Fig. 1E, top right), and use $\mathbf{x}_F$ as the input to the LGBM classifier under the same cross-validation (CV) protocol used earlier. This baseline combines information from both alignments directly, without explicitly enforcing consistency between paired SL and RL latent embeddings.

##### 3.3 Contrastive SL-RL (CSR) fusion

For CSR, building on the latent embeddings extracted from the SL and RL views, we employ a CLIP-based contrastive learning framework [1] to explicitly align paired representations in a shared embedding space (Fig. 1E, bottom right). Let $\mathbf{x}^{(i)}_{SL}$ and $\mathbf{x}^{(i)}_{RL}$ denote the latent representations of the same HFA sample under the SL and RL views, respectively. These are passed through modality-specific projection heads, denoted by $P_{SL}$ and $P_{RL}$, to obtain projected embeddings $\mathbf{z}^{(i)}_{SL} = P_{SL}(\mathbf{x}^{(i)}_{SL})$ and $\mathbf{z}^{(i)}_{RL} = P_{RL}(\mathbf{x}^{(i)}_{RL})$, which are subsequently $\ell_2$-normalized to yield unit embeddings $\mathbf{e}^{(i)}_{SL}$ and $\mathbf{e}^{(i)}_{RL}$. This design is motivated by prior contrastive learning studies showing that projection heads, normalised embeddings, and temperature scaling improve representation learning [2]. Given a minibatch of $B$ paired samples $\{(\mathbf{e}^{(i)}_{SL}, \mathbf{e}^{(i)}_{RL})\}_{i=1}^{B}$, we compute the cross-view similarity matrix with entries $s_{ij} = \mathbf{e}^{(i)\top}_{SL}\mathbf{e}^{(j)}_{RL}$, which correspond to cosine similarity due to $\ell_2$ normalization. Training increases similarity for matched SL-RL pairs while decreasing similarity for mismatched pairs within the minibatch using a symmetric contrastive cross-entropy objective:

$$\mathcal{L}_{CSR} = \frac{1}{2}\left(\mathcal{L}_{SL\to RL} + \mathcal{L}_{RL\to SL}\right) \tag{4}$$

where

$$\mathcal{L}_{SL\to RL} = -\frac{1}{B}\sum_{i=1}^{B}\log\frac{\exp(s_{ii}/\tau)}{\sum_{j=1}^{B}\exp(s_{ij}/\tau)} \tag{5}$$

and $\mathcal{L}_{RL\to SL}$ defined analogously. Here, $\tau$ denotes the temperature parameter that scales the logits before the softmax operation. This objective encourages the diagonal entries of the SL-RL similarity matrix, corresponding to correctly paired views from the same sample, to dominate over the off-diagonal entries corresponding to mismatched pairs. After contrastive training, we construct the classifier input as

$$\mathbf{e}_F = [\mathbf{e}_{SL}, \mathbf{e}_{RL}, \mathbf{e}_{SL} - \mathbf{e}_{RL}] \tag{6}$$

and use $\mathbf{e}_F$ for further classification under the same evaluation protocol. In this way, CSR not only preserves complementary information from the SL and RL views but also explicitly incorporates their aligned common structure and residual difference into a unified representation.

##### 3.4 Hyperparameter tuning for machine learning models

Hyperparameter tuning was conducted using RandomizedSearchCV for SL and RL separately, and the results of LGBM were analysed using SHAP values with different CV frameworks, including LOSO, SGKF, and SKF. In our experiments, for SL, the LGBM model used $num\_leaves = 70$ (maximum tree leaves), $max\_depth = 20$ (tree depth limit), $learning\_rate = 0.05$ (step size), $n\_estimators = 500$ (number of boosting rounds), $min\_data\_in\_leaf = 20$ (minimum samples per leaf), $min\_gain\_to\_split = 0.0$ (minimum gain to split), $feature\_fraction = 0.9$ (fraction of features sampled per iteration), $bagging\_fraction = 1.0$ and $bagging\_freq = 10$ (subsampled data every 10 rounds), $lambda\_l1 = 0.0$ (L1 regularization), $lambda\_l2 = 0.1$ (L2 regularization), objective = 'binary' (binary classification), $class\_weight$ = 'balanced' (handle class imbalance), $random\_state$ (reproducibility), $n\_jobs = -1$ (use all CPU cores), $device\_type$ = 'gpu' (GPU training), and $max\_bin = 255$ (histogram bins). For the RL, we switched to $bagging\_freq = 5$ (more frequent bagging), $lambda\_l1 = 0.1$, and $lambda\_l2 = 1.0$, retaining all other settings unchanged. For fusion (both ESR and CSR), we used $min\_data\_in\_leaf = 30$, $bagging\_freq = 5$, $lambda\_l1 = 0.5$, and $lambda\_l2 = 0.1$, retaining all other settings unchanged.
